# Homologs of the plastidal preprotein translocase Tic20 mediate organelle assembly in bacteria

**DOI:** 10.1101/2023.11.16.567405

**Authors:** Anja Paulus, Frederik Ahrens, Annika Schraut, Hannah Hofmann, Tim Schiller, Thomas Sura, Dörte Becher, René Uebe

## Abstract

Organelle-specific protein translocation systems are essential for organelle biogenesis and maintenance in eukaryotes but thought to be absent from prokaryotic organelles. Here, we identified that MamF-like proteins involved in the formation of bacterial magnetosome organelles share an ancient origin with Tic20 protein translocases found in chloroplasts. Deletion of *mamF*-like genes in the alphaproteobacterium *Magnetospirillum gryphiswaldense* results in severe defects in organelle positioning, biomineralization, and magnetic navigation. Consistent with translocase-like functions, these defects are caused by the loss of magnetosome targeting of a subset of organellar proteins containing C-terminal glycine-rich integral membrane domains. Our findings suggest that organelle-specific protein translocation systems may indeed play a role in bacterial organelle formation.

## Main Text

Eukaryotes possess macromolecular machineries like preprotein translocases of chloroplasts or mitochondria (Tic/Toc or Tim/Tom) to ensure biogenesis and maintenance of these organelles. Both translocases have a mixed phylogenetic origin and evolved through the combination of novel eukaryotic proteins with bacterial subunits of endosymbiotic origin (Muñoz-Gómez et al., 2015; Rast et al., 2015). Although the endosymbionts (cyano- and alphaproteobacteria, respectively) likely also owned membranous organelles, no organelle-specific protein import pathways equivalent to eukaryotic translocases have been identified among bacteria yet (Greening and Lithgow, 2020). Therefore, bacterial organelle biogenesis must proceed through distinct mechanisms. Magnetosomes, unique magnetic organelles, for example, emerge from the inner cell membrane by invagination (Komeili et al., 2006). In *Magnetospirillum gryphiswaldense* MSR-1 (MSR) and related alphaproteobacteria, this process is supposedly induced by assembly of membrane-embedded complexes of magnetosome proteins (MAPs) that initiate magnetosome membrane (MM) formation through a molecular crowding-like mechanism (Raschdorf et al., 2016). Notably, several of these MAPs also facilitate subsequent steps of magnetosome biogenesis like iron uptake, magnetite biomineralization, or crystal growth (Uebe and Schüler, 2016). Concurrently, a dynamic cytoskeletal network composed of the MM adaptor protein MamJ (Scheffel et al., 2006), which mediates magnetosome binding to the actin-like protein MamK (Komeili et al., 2006) and the membrane-bound mechanical scaffold MamY (Toro-Nahuelpan et al., 2019), orchestrates the assembly of approximately 30 magnetosomes into a linear magnetosome chain (MC). This arrangement produces a magnetic dipole that enables geomagnetic navigation. Although magnetosomes represent one of the best characterized bacterial organelles, the molecular mechanisms of magnetosome assembly and protein targeting are still poorly understood as it remained unknown how and when proteins are targeted to these organelles. Here, by combining bioinformatic in-depth analyses with systematic molecular, cell biological, quantitative proteomic as well as biochemical studies and evaluation of magnetism-related phenotypes, we show that magnetosomal MamF-like proteins (MFPs) and the distantly related plastidal preprotein translocase Tic20 are part of an ancient, widespread Tic20/HOTT (<u>H</u>omologs <u>o</u>f <u>T</u>ic <u>T</u>wenty, formerly DUF4870) superfamily that mediates organelle-specific protein targeting.

## Results

### MFPs are members of a common Tic20/HOTT superfamily

Recently, we discovered a ∼10 kb region within the genome of MSR that encodes several previously unrecognized MAPs (Uebe et al., 2018b). One of these proteins, MmxF, is closely related to the well-known MAPs MamF and MmsF (hereafter collectively termed MamF-like proteins, MFPs) that are thought to facilitate magnetite crystal growth by direct interaction with the crystal surface or metal ions (fig. S1A) (Murat et al., 2012; Lohße et al., 2014; Rawlings et al., 2014). Contrary to previous studies that failed to detect any homologs outside of magnetotactic bacteria, we identified highly significant homologies between MFPs, the DUF4870 (hereafter termed HOTT) as well as the Tic20 protein families (fig. S1B). Interestingly, while the widely distributed HOTT family encompasses only uncharacterized proteins, the Tic20 family, among uncharacterized cyanobacterial representatives, also comprises Tic20 from chloroplasts (fig. S1, C and D). In these photosynthetic organelles, Tic20 functions as a protein-translocating channel within the Tic complex located at the inner envelope membrane (IEM) (Kouranov et al., 1998; van Dooren et al., 2008; Kikuchi et al., 2009; Kovács-Bogdán et al., 2011). The identified homologies are further supported by cluster analysis of sequences (CLANS) (*17*). Here, the Tic20 family forms a widely separated but connected cluster to the HOTT family, in which MFPs seem to represent a discrete subfamily (Fig. 1A). Similar results were obtained by phylogenetic analyses where seven subfamilies including MFPs form distinct branches within the HOTT family, which itself is separated from the Tic20 family clade by a long branch (fig. S1E). Remarkably, despite only remote sequence homologies, structural superimposition, and multiple sequence alignments of MFPs, HOTT and Tic20 family proteins revealed similar architectures, including a reentrant helix, followed by a conserved charged loop and two additional transmembrane helices (Fig. 1B and C). The HOTT (including the MFPs) and Tic20 families thus form a common protein superfamily.

**Fig. 1.**
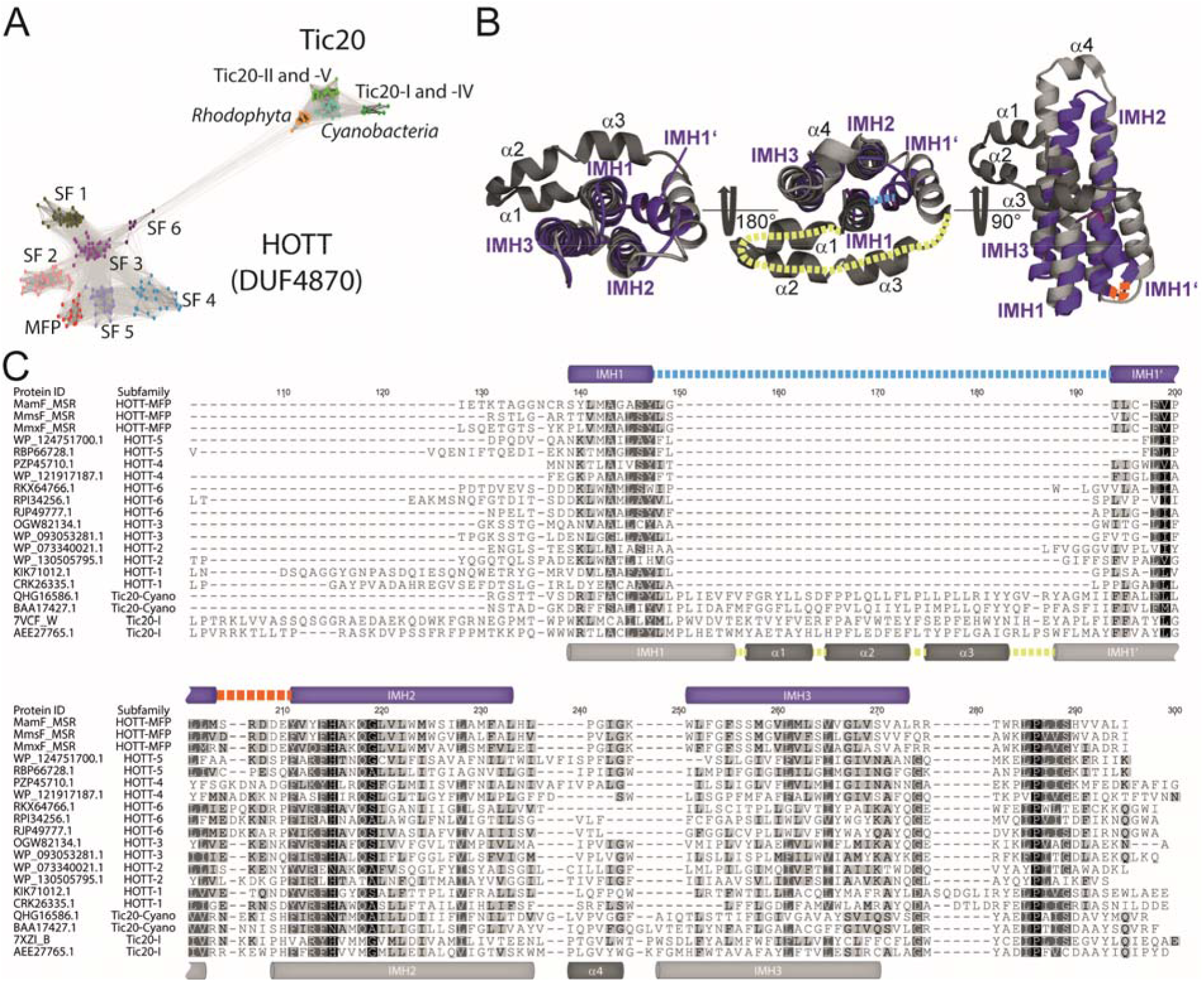
MFPs are members of a common Tic20/HOTT superfamily. **(A)** CLANS analysis of 229 TIC20/HOTT superfamily proteins. Protein sequences (colored dots) with greater pairwise similarity, are clustered closer together and connected by lines when pairwise BLAST P E-values are below 10^−5^. Accession numbers are listed in data table S2. **(B)** Superimposition (RMSD 3.419 Å, 61 to 61 atoms) of the Alpha fold prediction of the HOTT family member MamF (MSR-1, AF: Q6NE74, purple) (Jumper et al., 2021; Varadi et al., 2022) with the PDB-structure of Tic20 (*C. reinhardtii*, PDB: 7xZI, grey) (Liu et al., 2023) exhibits equal architectures including a reentrant-like helix, followed by a conserved charged loop (dashed orange line) and two additional transmembrane helices. All HOTT homologs have a short loop (dashed blue line), while Tic20 representatives contain an extended multi helical loop (dashed yellow line) in between the reentrant helix. RMSDs of superimpositions are listed in data table S2. **(C)** Multiple aa sequence alignment of highly conserved transmembrane regions of representatives of each subfamily highlights a shorter length of HOTT family members in comparison to the Tic20 family. Amino acid residues are shaded using the similarity color scheme and the Pam250 score matrix. Identical (100%), strongly (80-100%) or weakly (60-80%) similar residues are shaded in black, dark grey or light grey, respectively. Positions of predicted or definitive IHMs and alpha helices are indicated by purple (HOTT family) or grey (Tic20 family) cylinders. The reentrant-like helices of MamF (MSR-1) as well as Tic20 (*C. reinhardtii*) and the conserved acidic amino acids within loop 1 are marked with dashed lines, blue, yellow, and orange respectively. Accession numbers are listed in data table S2. (See also fig. S1 and data table S2)

### MFPs are essential for MC formation and magnetotaxis

The distant homology to Tic20 suggests that MFPs, besides regulating biomineralization, may also play a role during magnetosome assembly. To explore this hypothesis, we first analyzed transmission electron micrographs (TEM) of MSR mutants carrying deletions of *mamF*-like genes in all possible combinations, including the MFP-free mutant ΔF3 (Δ*mamF*Δ*mmsF*Δ*mmxF*) (Fig. 2 and fig. S2). These analyses revealed that, in agreement with known MFP deletion phenotypes (Murat et al., 2012; Lohße et al., 2014), the average magnetite crystal size gradually decreased from the WT, the single and double deletion mutants to the ΔF3 strain while crystal numbers per cell were only slightly reduced (Fig. 2, A and B). Beyond these expected biomineralization defects, however, the simultaneous deletion of all MFPs also resulted in the unexpected loss of MC formation as magnetosomes are randomly distributed within cells of strain ΔF3 (Fig. 2C). This observation is also reflected by an increased fraction of magnetosomes that have no neighboring particle within distance of 35nm and an overall lower number of close-neighboring magnetosomes (Fig. 2, D and E, fig. S2, B and C). Notably, expression of each *mamF*-like gene within ΔF3 restored magnetosome crystal sizes at least partially, whereas MC formation was reconstituted by expression of *mmsF* or *mmxF* but not *mamF* (Fig. 2, A and C, fig. S2A). Thus, only MmsF and MmxF seem to have an additional function during MC assembly.

**Fig. 2.**
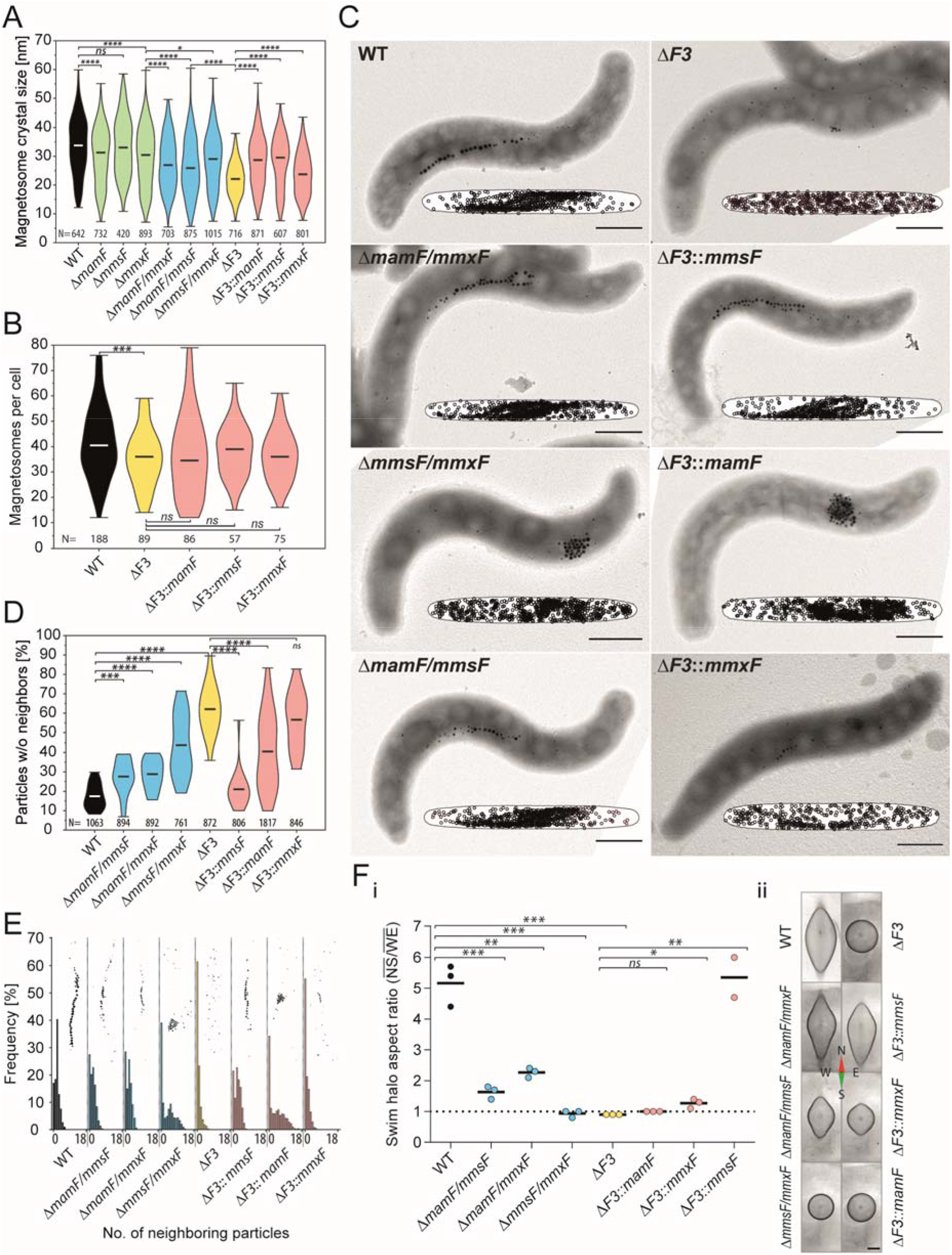
MFPs are essential for MC formation and magnetotaxis. **(A)** Deletion of *mamF*-like genes affects magnetite biomineralization and results in smaller crystal sizes. Violin plot displaying the magnetite crystal size distribution of the MSR WT (black), single (green), double (blue), and triple (ΔF3, yellow) MFP mutants as well as single MFP complemented ΔF3 strains (red). Particle diameters given in nm were measured from TEM micrographs. The number of analyzed particles [N] is indicated below each plot. The min, max, and mean values are indicated by bars. Statistical significance was estimated using an unpaired two-tailed Mann-Whitney U-test. Asterisks indicate the points of significance, *, P value ≤ 0.05; **, P value ≤ 0.01; ***, P value < 0.001; ****, P value < 0.0001; ns, not significant (P ≥ 0.05). Raw data is provided in data table S3. **(B)** Deletion of *mamF*-like genes has only a slight effect on particle numbers per cell. Violin plot showing the distribution of magnetosome numbers per cell in the WT, ΔF3, and the MFP complemented strains. Magnetosomes per cell were measured from TEM micrographs. The number of analyzed cells [N] are indicated. Coloring and statistical analysis like Fig. 2A. Raw data is provided in data table S3. **(C)** Magnetosome chain formation is disrupted in the absence of *mmsF* and *mmxF*. TEM micrographs of representative WT and various MFP mutant cells. Insets visualize normalized cellular distribution of magnetosomes (black circles) from at least 16 cells and 593 particles. (Scale bars: 500 nm) **(D)** Quantitative magnetosome neighbor analyses (qMNA; for details see fig. S2B and S2C) visualized in violin plots as the frequency of magnetosomes per cell that have no neighboring magnetosome within 35 nm around the outer edge of the magnetite crystal. Statistical significance was estimated using an unpaired two-tailed t-test with Welchs’ correction. The number of analyzed particles [N] measured from TEM micrographs are indicated. The min, max, and mean values are indicated by bars. Coloring and statistical significance similar to Fig. 2A. Raw data is provided in data table S3. **(E)** qMNA histograms of the WT, double deletion mutants, ΔF3 and single MFP complemented strains demonstrating the frequency of particle neighbor numbers within 35 nm around the outer edge of a magnetite crystal. Insets illustrate representative magnetosome particle distribution schemes. Sample size and coloring similar to Fig. 2A. Raw data is provided in data table S3. **(F)** MFPs are essential for magnetotaxis. i Swim halo aspect ratio of the WT and various MFP mutant strains within a homogenous 600 µT magnetic field 24 h after inoculation. Dots colored similar to Fig. 2A represent aspect ratios of individual biological replicates. Bars represents the mean. Statistical significance was estimated using an unpaired two-tailed t-test (labels similar to Fig. 2A). Raw data is provided in data table S3. ii Representative swim halos of the WT and various MFP mutant strains within a homogenous 600 µT magnetic field formed after 72 h. (Scale bar: 1 cm) (See also figure S2 and data table S3.)

Both, crystal size and MC formation are important parameters that influence the cellular magnetic dipole moment and, therefore, strongly affect the cells’ ability to align with the geomagnetic field (Pfeiffer and Schüler, 2020). Consequently, MFP mutants with reduced crystal sizes and defective MCs are severely impaired in magnetotaxis while strains exhibiting long MCs (WT and ΔF3::*mmsF*) showed strong magnetic alignment (Fig. 2F). Collectively, these results indicate that MFPs are essential for magnetic navigation and thus organellar function through regulation of MC formation and magnetosome crystal sizes.

### MFPs mediate MM protein assembly

The absence of MCs suggests that the function of the cytoskeletal proteins MamK, MamY, or MamJ might be affected in strain ΔF3. Nevertheless, these proteins are present at WT levels in ΔF3 cell extracts and even appear to be functional since fluorescently labelled proteins still localized in WT-like filaments in ΔF3 strains (Fig. 3, A and B). In purified MM extracts of ΔF3, however, the amounts of MamY and MamJ were significantly reduced and only MamK was detectable at WT levels (Fig. 3A). Since MM localization of MamY was previously shown to depend on MamJ (Toro-Nahuelpan et al., 2019), we surmised that loss of MamJ MM targeting is the primary cause for the absence of MCs in ΔF3. This suggestion is supported by decreased MamJ abundances in MM extracts of all MFP mutant strains that simultaneously lack *mmsF* and *mmxF* as well as the phenotypic similarity of the *mamJ* deletion mutant (Scheffel et al., 2006) with strains ΔF3::*mamF* and Δ*mmsF*Δ*mmxF*, respectively (fig. S2A, Fig. 3C). Consistently, quantitative proteomic analyses of WT and ΔF3 MM extracts revealed eleven proteins that are depleted in ΔF3 MM fractions. These include MamJ with a 44-fold and MamY with a 6-fold reduced abundance, while MamK was again identified in equal quantities in WT and ΔF3 MM extracts (Fig. 3D). Additionally, the magnetite crystal size-regulating proteins MamD, Mms5, and MamR (84-fold, 12-fold and 9-fold decrease, respectively) (Arakaki et al., 2014; Lohße et al., 2014) were also strongly depleted in ΔF3 MMs, which was verified by immunoblotting of isolated MM extracts of WT and ΔF3 strains expressing GFP fusions (Fig. 3D, fig. S3A). Interestingly, while MamF fails to mediate MamJ MM targeting, *in vivo* fluorescence imaging of WT and ΔF3 strains producing GFP-tagged MamD, Mms5, or MamR showed that expression of *mmsF*, *mmxF*, or *mamF* in ΔF3 individually restored magnetosome localization of MamD-, Mms5- and MamR-GFP, respectively (fig. S3B). These findings indicate that all MFPs are central for correct MM assembly but may differ in their “substrates”.

**Fig. 3.**
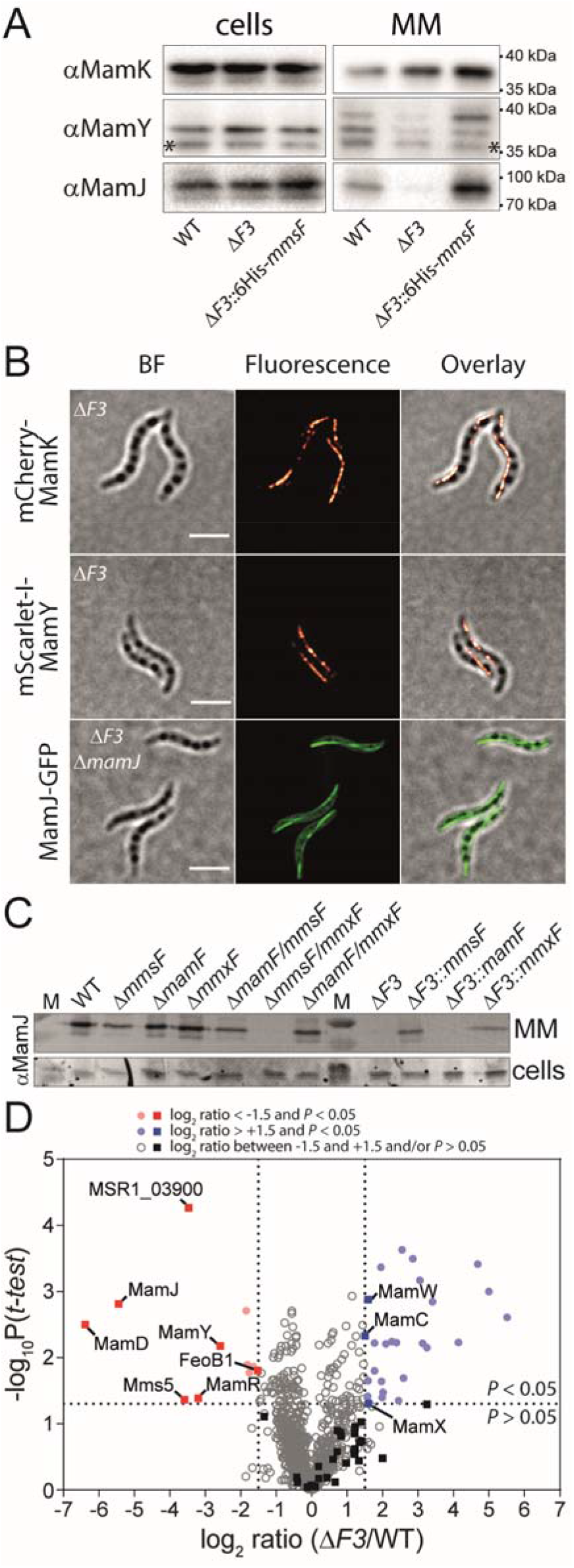
MFPs mediate MM protein assembly. **(A)** Immunodetection of MamK, MamY, and MamJ in whole cell extracts and isolated MM fractions of the WT, ΔF3, and ΔF3::*6His-mmsF* strains. Molecular weight standards are indicated. Unspecific bands are marked with asterisks. **(B)** Magnetosome-specific cytoskeletal proteins are still functional in ΔF3. Representative 3D-SIM micrographs showing WT-like filaments of fluorescently labelled MamK, MamY, and MamJ proteins in ΔF3 strains. BF, bright field image. (Scale bars: 2 µm) **(C)** Immunodetection of the MM adaptor protein MamJ in whole cell extracts and isolated MM fractions of the WT, single and double deletion mutants, the triple deletion mutant, and its single MFP complemented strains. MamJ, which is present in all cell extracts, is absent in MM fractions of all strains carrying simultaneous deletions of *mmsF* and *mmxF*. Electrophoretic mobility corresponds to a ∼90 kDa protein (*22*). **(D)** The protein composition of the MM in ΔF3 is strongly altered. Volcano plot showing the log2-fold change of proteins in purified magnetosome fractions of ΔF3 compared to the WT as determined by liquid chromatography–tandem mass spectrometry (LC-MS/MS). Magnetosome island (MAI)-encoded proteins (squares) with significantly altered abundance (log2 ratio <-1.5 or >1.5 and P < 0.05, vertical and horizontal dotted lines, respectively) compared to the WT are labeled. Circles represent non-MAI encoded proteins. Significantly depleted or enriched proteins are labelled with red and blue symbols, respectively. Non-significant proteins are depicted by grey or black symbols, respectively. Values are the mean of three biological replicates. (See also data table S4.) (See also figure S2A and S3, data table S4, and supplementary text)

### MamJ is embedded within the MM

To gain a better understanding of the MFP-mediated MM assembly, we chose to study MamJ as model “substrate” because of its strong depletion in the ΔF3 mutant and the easily detectable *mamJ* deletion phenotype (Fig. 3D, fig. S2A). Since MamJ-GFP has a cytosolic localization in the absence of other MAPs (ΔM05, fig. S4A) (Dziuba et al., 2020), we initially assumed that MamJ might be membrane-associated by binding to MmsF or MmxF. To assess this assumption, solubilized MM protein complexes were fractionated by size exclusion chromatography (SEC). To reduce complexity, we used MMs of strain ΔF3Δ*mms5*Δ*mamK*Δ*mamY*::*GFP*-*mmsF* in which, due to the deletion of all genes encoding MamJ interaction partners like MamK, MamY, and MFPs, merely a complemented, functional GFP-MmsF fusion protein (Murat et al., 2012) can interact with MamJ. In contrast to our expectations, MamJ and GFP-MmsF elution profiles overlapped only partially (Fig. 4A). Thus, only a small fraction of MM-bound MamJ might be directly associated with GFP-MmsF. To evaluate this possibility, solubilized magnetosomes of *GFP-mmsF* and *GFP-mamF* complemented ΔF3 mutant strains as well as the WT and Δ*mamJ* were subjected to co-immunoprecipitation (Co-IP). In these analyses, MamJ was only detected in Co-IP extracts of strain ΔF3::*GFP*-*mmsF* (Fig. 4B). Nevertheless, even in these extracts only minute amounts of MamJ were detectable as most MamJ remained in the unbound fraction. Thus, indeed only few molecules of MM-bound MamJ are in direct contact with MmsF. Next, to assess if the majority of magnetosome-bound MamJ molecules might instead be associated with other MAPs, carbonate extractions with isolated MMs from a Δ*mamK*Δ*mamY* deletion strain were performed. This treatment disrupts protein-protein interactions but leaves protein-lipid interactions intact (Fujiki, Y., Hubbard, A.L., Fowler, S., and Lazarow Paul B., 1982; Vögtle et al., 2017). Therefore, peripherally associated MamJ should be extractable from the MM. Interestingly, however, contrary to the well-known magnetosome-associated protein MamA (Taoka et al., 2006), which could be efficiently extracted from the MM, MamJ, similar to the integral MAP MamM (Barber-Zucker et al., 2016), remained bound to magnetosomes (Fig. 4C). While all tested proteins could be completely extracted from the MM by detergent treatments, even repeated and extended incubations with alkaline, acidic, or high salt buffer solutions failed to solubilize MamJ or MamM from the MM (fig. S4B). Collectively, these results indicate that magnetosome-bound MamJ behaves as an integral membrane protein.

**Fig. 4.**
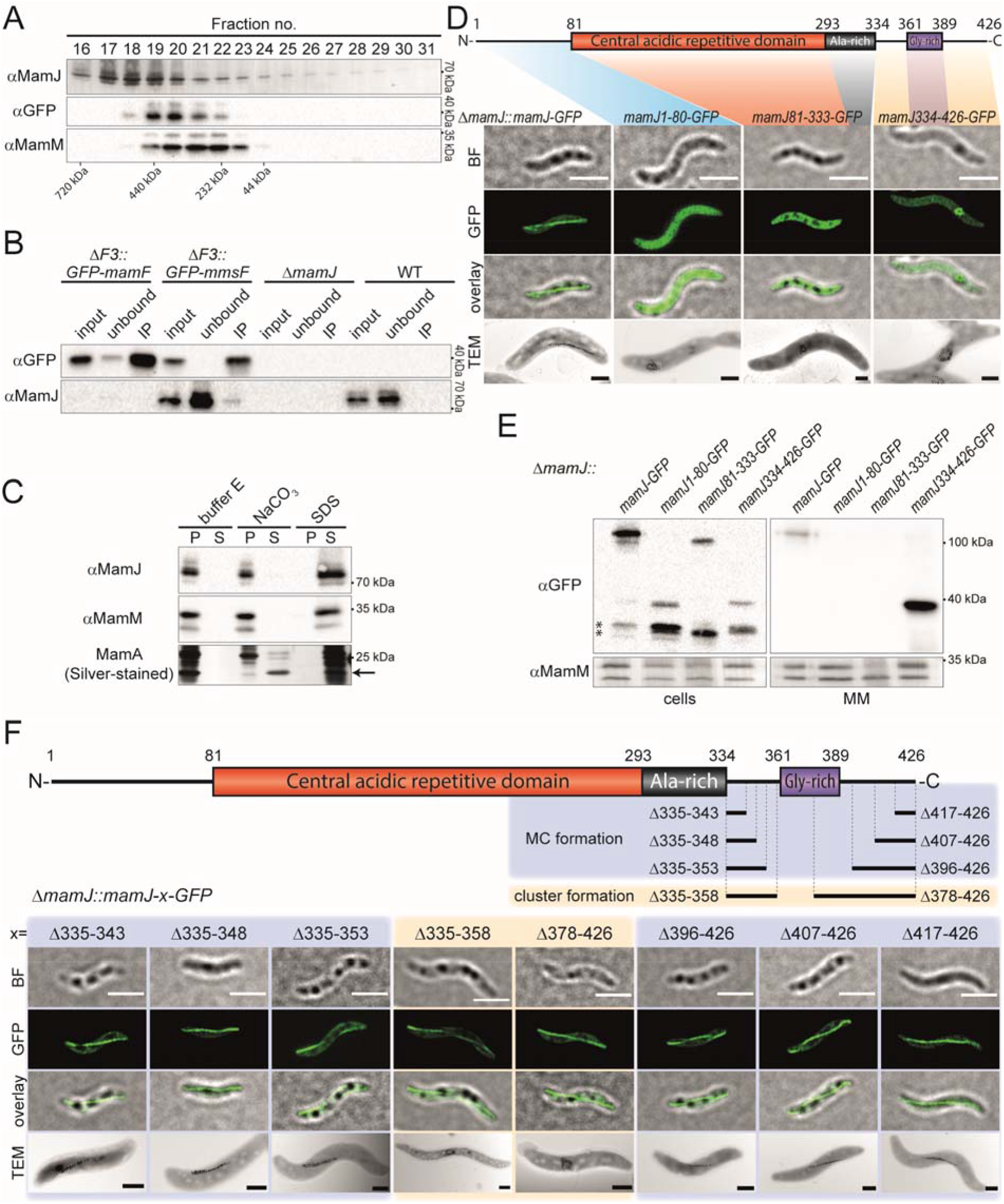
MamJ is inserted into the MM via its C-terminal domain. **(A)** Size exclusion chromatography (SEC) elution profiles of MamJ and GFP-MmsF overlap only partially. Preparative separation of solubilized MAP complexes of isolated magnetosomes from ΔF3DΔ*mamK*Δ*mamY*::*GFP*-*mmsF* using a Superose 6 increase 10/300 GL column. MAP complex elution fractions 16-31 were separated by SDS-PAGE and subsequently analyzed by Western blot immunodetection using MamJ, GFP, and MamM antibodies. Molecular weights of the SEC standard calibration are indicated. **(B)** Only minor amounts of magnetosome-bound MamJ directly interact with GFP-MmsF. Co-immunoprecipitation of solubilized magnetosomes from ΔF3::*GFP-mmsF*, ΔF3::*GFP-mamF*, WT and Δ*mamJ* mutant strains using anti-GFP nanobodies. Input, unbound, and immunoprecipitated (IP) fractions were analyzed by SDS-PAGE and subsequent Western immunoblots using GFP and MamJ antibodies. Molecular weight standards are indicated. **(C)** Magnetosome-bound MamJ behaves as an integral membrane protein. Western blot and SDS-PAGE analysis of buffer E-(negative control), carbonate-, and SDS-treated (positive control), magnetosome pellet (P) and supernatant (S) fractions from a Δ*mamKY* mutant show that MamJ and the integral MAP MamM are only detectable in supernatant fractions upon treatment with SDS. In contrast, carbonate efficiently extracted the peripheral and highly abundant MamA protein as shown by silver-stained SDS PAGE (black arrow). Molecular weight standards are indicated. **(D)** Domain structure of MamJ containing a central acidic repetitive (CAR) (red), alanine- and glycine-rich domain (black and purple, respectively). TEM and 3D-SIM imaging of full-length (aa 1-426) or truncated versions of MamJ fused to GFP and expressed in ΔmamJ reveal that only full-length MamJ-GFP is functional to restore the MCs. BF, bright field image. (Scale bars: SIM 2 µm, TEM 0.5 µm) **(E)** The C-terminal domain mediates magnetosome targeting of MamJ. Cell extracts and isolated magnetosomes of Δ*mamJ* strains producing different MamJ truncations fused to GFP were analyzed by immunoblotting using GFP antibodies. A Western blot against MamM is shown as loading control. Molecular weight standards are indicated, and unspecific bands are marked with asterisks. **(F)** The putative C-terminal IMH of MamJ is essential for MC reconstitution in Δ*mamJ*. TEM and 3D-SIM imaging of truncated MamJ-GFP variants produced in Δ*mamJ*. Domain structure is depicted analogue to Figure 4D. MamJ variants that maintained or lost the ability to reconstitute MC formation are colored in blue and apricot, respectively. BF, bright field image. (Scale bars: SIM 2 µm, TEM 0.5 µm) (See also figure S4)

### A hydrophobic C-terminus is required for MM targeting of MamJ

To investigate which domain of MamJ is required for its MM targeting, alleles encoding full-length (aa 1-426) or truncated *mamJ*-*GFP*-fusions were expressed in Δ*mamJ*. As shown previously (Scheffel et al., 2006), full-length MamJ-GFP restored MC formation and localized in a linear pole-to-pole spanning signal. Contrary, GFP fusions to the MamJ N-terminal (aa 1-80) or the alanine-rich central acidic repetitive (CAR, aa 81-333) domains revealed only soluble localization signals (Fig. 4D). Consequently, Δ*mamJ*-like magnetosome clusters were still observable by TEM. When GFP was fused to the MamJ C-terminus (aa 334-426) large fluorescent foci reminiscent of magnetosome clusters could be observed. Since a MamJ334-426-mCherry fusion colocalized with signals from a GFP fusion to the essential MAP MamB (Uebe et al., 2011) in large fluorescent foci within Δ*mamJ*, the MamJ C-terminus seems to mediate MamJ MM targeting but lacks the ability to bind the cytoskeleton (fig. S4C). Finally, immunoblotting of isolated MMs from complemented Δ*mamJ* strains confirmed that only full-length MamJ and the MamJ334-426-GFP fusion protein were targeted to the MM (Fig. 4E). Consistent with its ability to mediate MM targeting, the MamJ C-terminus interacts with MmsF and MmxF, but not MamF, in bacterial two-hybrid (BACTH) assays (fig. S4D) and encompasses a putative integral membrane helix (IMH, aa 359-379) (Scheffel and Schüler, 2007). Moreover, this glycine-rich (37.9% glycine content) region proved to be essential for MC formation as only truncations within or in close proximity of the IMH (Δ335-358, Δ378-426) prevented reconstitution of MC formation in Δ*mamJ* although all variants retained their ability to bind the cytoskeleton (Fig. 4F). These results indicate that MamJ is inserted into the MM via its hydrophobic, glycine-rich C-terminus. Notably, also MamD and Mms5 contain C-terminal IMHs of high glycine content, which are preceded by alanine-rich domains. Thus, the three most strongly affected proteins of the ΔF3 mutant share a similar domain architecture. Moreover, despite only low sequence similarity with MamJ, the C-terminal domains of MamD and Mms5 do also interact with each MFP in BACTH assays (fig. S4E). Thus, in agreement with our fluorescence microscopy results, all three MFPs facilitate MM targeting of MamD and Mms5.

### MFPs indirectly facilitate magnetite biomineralization

Previous studies suggested that MFPs directly promote magnetite crystal growth through interaction of iron ions with a cluster of conserved acidic amino acids within loop 1 (Fig. 1C) (Rawlings et al., 2014). Our findings, however, suggest that mistargeting of crystal growth-promoting proteins like MamD or Mms5 is the main contributor for decreased magnetite crystal sizes in MFP mutants. To discriminate between both possibilities, MmsF variants, lacking all acidic (D34N, D36N, D37N, E38N) or charged (D34N, R35Q, D36N, D37N, E38N) residues of loop 1 were tested for their ability to increase magnetosome crystal sizes in ΔF3 (Fig. 5A). Consistent with an indirect role during magnetite growth, both tested variants restored crystal sizes to the same level as wild-type MmsF (Fig. 5C). Next, we examined if MmsF maintains its ability to improve crystal growth when most of the proteins mistargeted in ΔF3 are absent. To this end, we deleted *mamR* within the ΔA13Δ*mms5*Δ*mmxF* mutant (Zwiener et al., 2021) that already lacks all MFPs and most MFP-dependent proteins due to the combined deletion of several accessory magnetosome gene operons (*mamGFDC*, *mms5*, *mms6*, and *mamXY*). Since all essential genes of the *mamAB* operon are maintained, the resulting ΔA13Δ*mms5*Δ*mmxF*Δ*mamR* mutant still formed tiny magnetite particles that are, because of the absence of MFPs, dispersed throughout the cell (Fig. 5B, data table S4). Upon complementation with *mmsF*, the ΔA13Δ*mms5*Δ*mmxF*Δ*mamR* mutant regained the ability to assemble MCs but crystal sizes showed only a minor increase compared to the expression of *mmsF* in the ΔF3 mutant (Fig. 5, B and C, data table S4). Thus, even though MmsF is functional in the ΔA13Δ*mms5*Δ*mmxF*Δ*mamR* mutant, as evidenced by MC restoration, the absence of MFP-targeted proteins prevented considerable magnetite crystal growth. In summary, our analyses indicate that MFPs indirectly regulate organelle positioning and magnetosome crystal size by mediating correct organelle assembly.

**Fig. 5.**
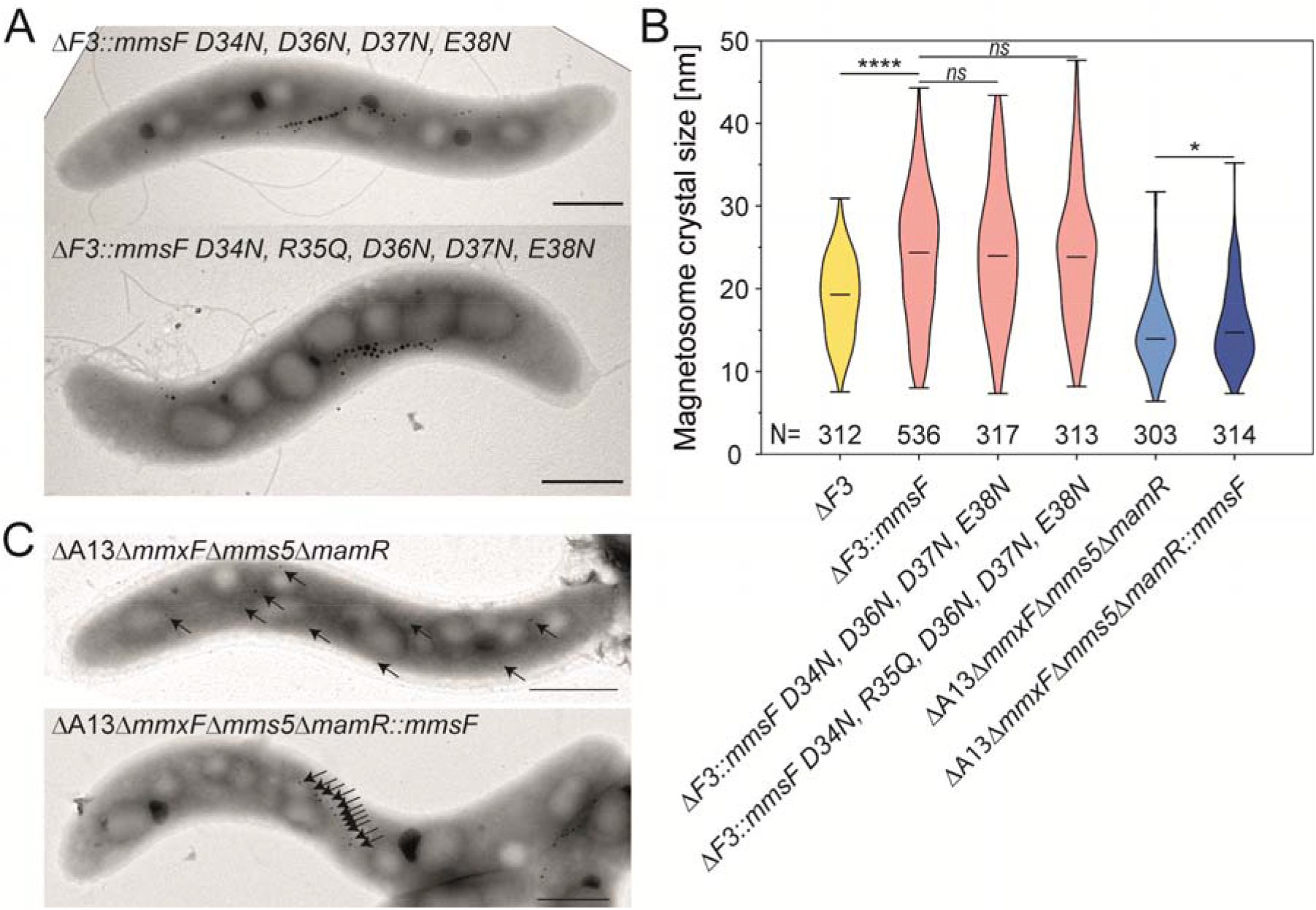
MFPs have an indirect effect on magnetite biomineralization. **(A)** Representative TEM micrographs of ΔF3 expressing *mmsF* mutant alleles lacking acidic (D34N, D36N, D37N, E38N) or charged (D34N, R35Q, D36N, D37N, E38N) residues within the loop between IMH1 and IMH2 (see Figure 1 B and C). (Scale bars: 500 nm) **(B)** Representative TEM micrographs of the ΔA13Δ*mms5*/*mmxF*/*mamR* mutant and the complemented strain ΔA13Δ*mms5*/*mmxF*/*mamR*::*mmsF*. The positions of the small magnetosomes are indicated by arrows. (Scale bars: 500 nm) **(C)** Violin plot showing the magnetite crystal size distribution of strain ΔF3 expressing WT and mutant *mmsF* variants as well as strains ΔA13Δ*mms5*/*mmxF*/*mamR* and ΔA13Δ*mms5*/*mmxF*/*mamR*::*mmsF*. The number of analyzed particles [N] is indicated. The min, max, and mean values are given by bars. Statistical significance was estimated using an unpaired two-tailed Mann-Whitney U test (*, P value ≤ 0.05; **, P value ≤ 0.01; ***, P value < 0.001; ****, P value < 0.0001; ns, not significant (P ≥ 0.05)). Raw data is provided in data table S5. (See also Figure 1 and data table S5)

## Discussion

In this study, we showed that MFPs of magnetotactic bacteria are members of the yet uncharacterized HOTT (DUF4870) protein family, which itself constitutes an ancient superfamily with Tic20 preprotein translocases (Fig. 1, fig. S1). Intriguingly, despite only distant sequence homology, plastidal Tic20 and bacterial MFPs share similar structures and play essential roles in proper organelle assembly and function. Deletion of all *mamF*-like genes in MSR, for example, resulted in the mislocalization of several MAPs, leading to magnetosome mispositioning, reduced magnetite biomineralization as well as loss of magnetotaxis (Fig. 2). Likewise, *Tic20-I* mutants of *A. thaliana* demonstrated impaired plastidal import of photosynthesis-related preproteins, resulting in abnormal chloroplast development and albino phenotypes (Kikuchi et al., 2009; Kasmati et al., 2011). Thus, in both cases only a small subset of organellar proteins is affected.

Tic20 mediates translocation of proteins across the inner plastidal envelop membrane (Kikuchi et al., 2013), while MFPs seem to insert proteins into the magnetosome membrane. The putative substrate MamJ, for instance, is soluble in the absence of MFPs but tightly binds to the MM in their presence, thus requiring detergents for extraction (Fig. 3C, fig. S4, A and B). Moreover, membrane binding seems to be independent of protein-protein interactions as magnetosome-bound MamJ is resistant against carbonate, high salt, acidic, or alkaline treatments and only a minor MamJ fraction interacts with MmsF (Fig. 4C, fig. S4B, Fig 3, A and B). Also consistent with a membrane insertion, MamJ functionality and magnetosome targeting depends on the presence and integrity of a putative C-terminal IMH (Fig. 4E and F) which represents one of the very few conserved regions within this hypervariable protein (Dziuba et al., 2023). Interestingly, two further potential MFP substrates, MamD and Mms5, share several of these characteristics with MamJ. Both proteins can also only be extracted from magnetosomes by use of detergents (Arakaki et al., 2003) and possess a soluble, alanine-rich N-terminal domain followed by a moderately hydrophobic (grand average hydropathy scores 1.3-1.6, (Peschke et al., 2019), glycine-rich C-terminal domain that interacts with MFPs. While the Ala-rich domains lack any sequence homology, the C-terminal domains share an IMH with a GxxxG glycine zipper motif that is commonly found within membrane proteins (Kim et al., 2005). Notably, such glycine zipper motifs are specifically enriched among chloroplast inner envelope membrane proteins that are inserted via a stop-transfer mechanism (Viana et al., 2010; Froehlich and Keegstra, 2011). However, the GxxxG motif is unlikely to serve as a universal recognition motif for MFPs as MamG and Mms6 harbor similar motifs but do not interact with MFPs and remain largely unaffected by their deletion (fig. S3, A and B, fig. S4, D and E). Thus, further studies are required to determine how MFPs recognize their substrates. In the future, this knowledge could be exploited to sustainably produce tailored, multifunctional magnetosomes for use in biotechnology and biomedicine (Rosenfeldt et al., 2021).

Additionally, our study reassigns the role of MFPs during magnetosome formation. While previous studies assumed an active crystal growth-promoting activity (Murat et al., 2012; Rawlings et al., 2014) our data show that MFPs are central for MM protein assembly and fail to significantly improve crystal growth in the absence of their substrates (Fig. 5). Their contribution to magnetite crystal maturation thus represents only an indirect effect that is based on the direct MM-targeting of proteins which themselves mediate crystal growth (e.g., MamD, Mms5).

Initially, we were surprised to identify homologs of the plastidal biogenesis factor Tic20 within an alphaproteobacterium that is distantly related to the ancestor of mitochondria. However, in contrast to Tic20, whose occurrence is restricted to plastid-containing eukaryotic linages (11 phyla) and cyanobacteria, the homologous HOTT family is widely spread among all domains of life (present in at least 38 bacterial, 13 archaeal, and 5 eukaryotic phyla). This extensive phylogenetic distribution and the larger number of divergent subfamilies suggest that the HOTT family is evolutionarily more ancient. Thus, although the short protein lengths, long evolutionary distances, and high sequence divergences among superfamily representatives (van Dooren et al., 2008; Machettira et al., 2011) prevented more precise phylogenetic reconstructions (fig. S1C and D), our findings support the hypothesis that Tic20 originated from a HOTT ancestor that probably already mediated protein targeting. In contrast, previous studies suggested that Tic20, similar to the mitochondrial inner membrane preprotein translocases (Tim17/22/23), evolved from bacterial LivH-like amino acid transporters (Rassow et al., 1999; Reumann, 1999; Bodył et al., 2010). In support of our data, however, several subsequent studies failed to detect any significant homologies between LivH, Tic20, or Tim17/22/23, respectively (Gross and Bhattacharya, 2009; Kasmati et al., 2011; Žárský and Doležal, 2016).

In summary, we not only present the first functional characterization of the yet uncharacterized HOTT protein family but also provide insights into its evolutionary relationship and functional conservation with the Tic20 protein family. Therefore, our findings also represent the first evidence for at least primitive organelle-specific translocases within bacteria and shed new light on the evolution of eukaryotic organelles.

## Acknowledgments

We would like to thank Gabriele Zaus and Ulrike Brandauer for technical assistance as well as Stephanie Bauer and Michael Gass for help during immunoblot analysis, two-hybrid assays, strain, and plasmid construction.

## Funding

The authors are grateful for financial support by the DFG, German Research Foundation grant UE200/1-1 (RU)

DFG, German Research Foundation grant INST 91/374-1 LAGG (RU)

Further funding was received from the BMBF, Federal Ministry of Education and Research grant MagBioFab (RU)

## Author contributions

Conceptualization and methodology: RU

Investigation: AP, FA, AS, TSc, HH and RU (conducted the biochemical and cell biological studies), TS and DB (performed MS proteomics and analyzed MS data)

Formal analysis: RU

Funding acquisition: RU

Supervision: RU

Writing - Original draft: AP and RU

Writing - Review & editing: all authors.

## Competing interests

Authors declare that they have no competing interests.

## Data and materials availability

All data supporting the findings of this study are included in the main text, the supplementary materials or the auxiliary data files. The mass spectrometry proteomics data have been deposited to the ProteomeXchange (http://proteomecentral.proteomexchange.org) consortium via the PRIDE partner repository (Perez-Riverol et al., 2022) with the dataset identifier PXD032959

## Supplementary Material

### Materials and Methods

#### Bacterial strains and culture conditions

All strains used in this study are listed in table S1. Strains of *Escherichia coli* were grown in lysogeny broth (LB) supplemented with kanamycin (25 µg mL^-1^) or ampicillin (50 µg mL^-1^) at 37 °C (except BTH101 which was grown at 28 °C) and shaking at 170 rpm. For cultivation of *E. coli* WM3064, 0.1 mM DL-α,ε-diaminopimelic acid (DAP) was added. Unless stated otherwise, *Magnetospirillum gryphiswaldense* strains were grown micro aerobically at 28 °C in modified flask standard medium (FSM) with moderate agitation (120 rpm) (Heyen and Schüler, 2003). For selection of chromosomal insertions, tetracycline (20 µg mL^-1^), chloramphenicol (5 µg mL^-1^) or kanamycin (5 µg mL^-1^) were added.

#### Molecular and genetic techniques

Molecular techniques were performed using standard protocols (Sambrook and Russell, 2001). All constructs were sequenced by Macrogen Europe (Amsterdam, Netherlands).

#### Plasmids and primers

All plasmids and primers used in this study are listed in table S2 and table S3, respectively.

#### Generation of site-specific chromosomal insertion and deletion mutants

For the generation of unmarked deletion mutants, ∼0.7 kb fragments of the up- and downstream flanking regions of the corresponding gene were amplified by PCR using Phusion polymerase (NEB) and primers. After gel purification of PCR products, the corresponding up- and downstream fragments were fused by overlap extension PCR using T4 polynucleotide kinase (Thermo Scientific) phosphorylated primers. Fused PCR products were then cloned into an EcoRV-linearized and dephosphorylated pORFM-GalK-MCS deletion vector. Plasmids were subsequently transferred to MSR by conjugation as described previously (Barber-Zucker et al., 2016). After incubation for 5 days (28 °C, 2% O_2_), kanamycin-resistant plasmid insertion mutants were transferred to 100 µL fresh FSM, and grown overnight (28 °C, 2% O_2_). 100 µl cultures were then inoculated into 900 µL fresh FSM and incubated for 24 h at 28 °C and 2% O_2_ before 200 µL were plated on FSM agar plates containing 2.5% (w/v) galactose and 100 ng mL^-1^ anhydrotetracycline. After incubation for additional 5 days (28 °C, 2% O_2_), mutant colonies were transferred to 100 µL FSM and verified by PCR.

#### Construction of MSR expression vectors

For complementation of *mamF*-like mutant strains, *mamF*, *mmsF*, and *mmxF* were PCR-amplified with specific primers, digested with KpnI/SacI, and cloned into similarly digested pBAM-Tet-*mamD*-GFP vector (Uebe et al., 2018a) to replace *mamD*-*GFP* against *mamF*-like genes. The resulting constructs were then transferred to MSR mutants via conjugation. For expression of *mamF*-like-*GFP* fusions, *GFP* together with a helix linker was amplified by PCR from pBAM160-*GFP*-*ccfM* (Pfeiffer et al., 2020), digested with KpnI and cloned into KpnI-digested pBAM-Tet-*mmsF* and pBAM-Tet-*mamF* vectors, respectively. The resulting plasmids pBAM-Tet-*GFP*-*mmsF* and pBAM-Tet-*GFP*-*mamF* were then transferred to MSR strain via conjugation.

For expression of additional fluorescently labeled proteins in MSR, *mamR*, *mms5*, and *mamG* were PCR-amplified with specific primers (table S3), digested with KpnI/XbaI, and cloned into the similarly digested pBAM-Tet-*mamD*-*GFP* vector (Uebe et al., 2018a) to replace *mamD*. For co-expression with *mamF*-like genes, *mamD*-*GFP*, *mamG*-*GFP*, *mamR*-*GFP*, and *mms5*-*GFP* containing pBAM-Tet vectors were digested with EcoRI/SacI to cut out the complete expression cassette including the constitutive *mamG* promotor. The resulting DNA fragments were then cloned into EcoRI/SacI-digested pBAM1 (Martínez-García et al., 2011) vector containing a kanamycin resistance gene. These constructs were then transferred to MSR mutants via conjugation. For co-expression, transfers were performed successively in independent conjugation experiments.

For expression of *mCherry*-*mamK*, the *mCherry*-*mamK* construct was amplified from MSR::*mCherry*-*mamK* containing a chromosomal in frame fusion (Raschdorf et al., 2014). The PCR product was digested with NdeI/XbaI and ligated into NdeI/XbaI-digested pBAM-P_tet_-*mamA-GFP* for exchange against *mamA*-*GFP* (Pfeiffer et al., 2020). Similarly, a synthetic *mScarlet-I*-*mamY* (ATG:biosynthetics GmbH, Merzhausen, Germany) was inserted into the NdeI/XbaI restrictions sites of pBAM-P_tet_-*mamA*-GFP to yield pBAM-P*_tet_*-*mScarlet-I*-*mamY*.

For the expression of truncated *mamJ*-*GFP* fusions, selected regions were PCR-amplified and cloned into the NdeI/KpnI restriction sites of the vector pBAM-Tet-*mamJ*-*GFP* (Kolinko et al., 2014). Alternatively, an inverse PCR-based cloning strategy was used to generate C-terminal truncations.

To study the colocalization of the MamJ C-terminus with MamB-GFP, a *mamJ*334-426-*mCherry* fusion was generated. Therefore, the vector pBAM2-Tn7-Tet-P_lac_ (Chevrier et al., 2022) as digested with NotI/XhoI. The P*_lac_*-*mamB*_*lacI* containing fragment was then ligated into the NotI/XhoI sites of pBAM2-Tn7-Cam (unpublished). Subsequently, a PCR-generated *mamJ*334-426-mCherry fusion was inserted into the NdeI/NotI restrictions sites to exchange *mamB* and yield pBAM2-Tn7-Cam-P_lac_-*mamJ*334-426-mCherry.

All constructs were verified by PCR and DNA sequencing.

#### Motility soft agar assay

Soft agar (0.2% agar (w/v)) swimming assays were conducted as described previously (Pfeiffer and Schüler, 2020), using modified FSM with a reduced concentration of potassium lactate (1.5 mM). Precultures were grown in triplicates per strain for at least three passages prior to the experiment in 6 well plates in FSM at 28 °C under defined microoxic condition (2% O_2_). Five microliters of cell suspension (adjusted to an OD_565_ of 0.1) were pipetted into 7 mL soft agar within 6-well plates and incubated at 28 °C under ambient oxygen conditions within a homogenous magnetic field of 0.6 mT using a custom coil setup. The plates were documented after one and two days of incubation and the swimming expansion was measured.

#### Bacterial two-hybrid assays

For protein interaction studies using the adenylate cyclase-based bacterial two-hybrid assay (Karimova et al., 1998), genes of interest were amplified from isolated chromosomal DNA of MSR and cloned into pUT18C, pUT18, pKT25, and pKNT25 plasmids. Therefore, the respective primers were phosphorylated before amplification using T4 polynucleotide kinase (Thermo scientific). Phosphorylated PCR-fragments were then ligated with SmaI-linearized and dephosphorylated (FastAP, Thermo scientific) vectors. The resulting T18- and T25-based plasmids were co-transformed into the *E. coli* BTH101 reporter strain. Cells were plated on LB agar supplemented with 40 µg mL^-1^ 5-bromo-4-chloro-3-indolyl β D galactopyranoside (X-Gal), 0.5 mM isopropyl β-D-1-thiogalactopyranoside (IPTG), ampicillin (100 µg mL^-1^) and kanamycin (50 µg mL^-1^) and incubated at 28 °C for 24 h. Finally, several colonies per plasmid combination were grown overnight at 28 °C in LB medium supplemented with IPTG (0.5 mM), ampicillin (100 µg mL^-1^) and kanamycin (50 µg mL^-1^), and 3 µL of culture were spotted onto M63 mineral salts agar containing 0.2% (w/v) maltose, X-Gal (40 µg mL^-1^), 0.5 mM IPTG, ampicillin (50 µg mL^-1^), and kanamycin (25 µg mL^-1^) (Uebe et al., 2019). M63 plates were then incubated at 28 °C for one day and documented with a Lumix FZ38 camera (Panasonic, Kadoma, Japan). Solely combinations with homogeneous, intense blue color formation were regarded as positive. Co-transformants harboring empty vectors or combinations of empty and test vectors of the respective T18- and T25-protein fusions served as negative controls on the same plates. All experiments were carried out in triplicates.

#### Magnetosome isolation

For magnetosome isolation, MSR strains were cultivated for 24 h at 28 °C and 120 rpm in 5 or 10 L flasks filled with 3 or 5 L FSM, respectively. For comparison of the magnetosome membrane proteome, MSR strains were cultivated by anaerobic fed-batch fermentation in 3 L Eppendorf bioreactors using a BioFlo 320 system (Eppendorf, Hamburg, Germany) (Lohße et al., 2016; Riese et al., 2020). To regulate the pH, 1 M KOH or an acidic feed composed of 1 M HNO_3_, 0.85 M potassium lactate, 0.45 M sodium nitrate, 25 mM magnesium sulfate and 3 mM Fe(III)-nitrate was pumped into the bioreactor when the pH changed by 0.1 from the set point of pH 7.0. After 30-35 h, cells were harvested by centrifugation at 8000 x g at 4 °C for 15 minutes (Avanti J-26 XP centrifuge; JLA 8.1000 rotor, Beckman Coulter, Brea, USA). Pellets were washed with buffer W (20 mM 4-(2-hydroxyethyl)-1-piperazineethanesulfonic acid (HEPES), 5 mM ethylenediaminetetraacetic acid (EDTA), pH 7.4) twice, pelleted by centrifugation, and stored at −20 °C until further use.

Magnetosome isolation was performed with an optimized protocol described recently (Rosenfeldt et al., 2021). Briefly, cell pellets were resuspended in ∼35 mL g^-1^ in buffer R (50 mM HEPES, 1 mM EDTA, 0.1 mM phenylmethylsulfonyl fluoride (PMSF), pH 7.4) and lysed by three cycles in a M110-L Microfluidizer processor (Microfluidics Corp., Westwood, USA) equipped with a H10Z interaction chamber at 1241 bar. Lysed cells were cleared from cell debris by centrifugation at 750 x g at 4 °C for 10 minutes (Allegra X-15R centrifuge; SX4750A rotor, Beckman Coulter, Brea, USA). The supernatant was loaded onto a magnetized MACS CS column (Miltenyi Biotec, Bergisch Gladbach, Germany) equilibrated in 50 mL of buffer E (10 mM HEPES, 1 mM EDTA, pH 7.4). The column was washed with 50 mL of buffer E, followed by 50 mL of buffer S (10 mM HEPES, 1 mM EDTA, 200 mM NaCl, pH 7.4) and again 50 mL of buffer E. Afterwards, the magnets were removed from the column and the magnetic fraction was eluted with 30 mL of ddH_2_O. Afterwards, HEPES and EDTA were added to a concentration of 10 mM and 1 mM, respectively, and 15 mL of this magnetic fraction was centrifuged on top of a 7 mL 60% (w/w) sucrose cushion in buffer E at 4 °C, 183,700 x g for 1.5 hours (Sorvall WX Ultra 80, Thermo Fisher Scientific, Waltham, MA, USA) (Raschdorf et al., 2018). The supernatant was discarded and the magnetosome pellet resuspended in 200 μL buffer E.

#### Protein concentration determination and normalization of magnetosome solutions

Protein concentrations were determined with the Roti-Quant universal kit according to the manufacturer’s protocol in flat-bottomed 96-well plates using an Infinite M200Pro plate reader (Tecan Group Ltd., Männedorf, Switzerland).

#### Gel electrophoresis and Western immunoblot

Analysis of proteins by denaturing sodium dodecyl sulfate polyacrylamide gel electrophoresis (SDS-PAGE) was performed on 8×10 cm or 10.5×10 cm acrylamide gels with 1.5 mm thickness at 300 V and 25 mA per gel (Hoefer SE250/SE260 chamber, Pharmacia Biotech, Upsala, Sweden; MP-300V power supply, Major Science, Saratoga, USA). If not stated otherwise, samples corresponding to 10 µg protein were mixed with 5x SDS-sample buffer (10% (w/v) SDS, 25% (v/v) β-mercaptoethanol, 25% (v/v) glycerol, 0.05% (w/v) bromophenol blue, 0.3 M Tris/HCl pH 6.8) to a 1-fold concentration and incubated at 95 °C for 10 minutes (ThermoMixer F1.5, Eppendorf, Hamburg, Germany) prior loading to the gels. For standard SDS-PAGE, 8-22.5% or 12-22.5% acrylamide linear gradient gels were used. In order to visualize protein bands, gels were stained with Coomassie brilliant blue R-250. To this end, gels were incubated for 30 minutes in Coomassie staining solution (50% (v/v) methanol, 10% (v/v) acetic acid, 0.1% (w/v) Coomassie R-250) followed by incubation in washing solution (10% (v/v) methanol, 7% (v/v) acetic acid) over-night. Alternatively, an improved silver staining method was used (Blum et al., 1987). To this end, gels were incubated in fixing solution (40% (v/v) methanol, 36.5% (v/v) of 37% (w/v) formaldehyde) for 10 minutes, washed twice in ddH_2_O for 5 minutes, incubated with 0.02% (w/v) thiosulfate solution for 1 minute before washing with ddH_2_O for 20 s twice. The gel was then covered with impregnating solution (0.1% (w/v) AgNO_3_) for 10 minutes in the dark. After shortly washing in ddH_2_O, a small volume of developing solution (3% (w/v) Na_2_CO_3_, 0.135% (v/v) of 37% (w/v) formaldehyde, 0.02% (v/v) of 0.02% (w/v) Na_2_S_2_O_3_) was used for washing the gel and then removed. The gel was incubated with fresh developing solution until protein bands became visible (1 to 5 minutes). The development was stopped by discarding the developing solution and incubating the gel in stopping solution (1.86% (w/v) EDTA) for 10 minutes. Stained gels were documented in ChemiDoc XRS+ (Bio Rad Laboratories Inc., Herkules, USA) or a DMC-FZ38 digital camera (Panasonic Corp., Kadoma, Japan).

Western blotting and immunodetection were performed according to protocols described previously (Dziuba et al., 2020).

#### Protein complex purification

For analysis of magnetosome protein complexes by size exclusion chromatography, isolated magnetosomes of MSR ΔF3Δ*mms5*Δ*mamK*Δ*mamY*::*GFP*-*mmsF* were solubilized using 0.1% (w/v) lauryl maltose neopentyl glycol (LMNG). To this end, isolated magnetosomes were adjusted to an OD_492_ of 6 (corresponding to 0.28 µg mL^-1^ protein) in buffer E to a final volume of 10 mL and pelleted by centrifugation at 21,130 x g and 4 °C for 30 minutes (Eppendorf 5424 R, FA-45-24-11 rotor). The pellet was resuspended in 2.5 mL of buffer E containing 50 mM NaCl, 0.1% (w/v) LMNG and incubated over-night at 4 °C under constant agitation. Subsequently, magnetosomes were removed from this solution by centrifugation at 21,130 x g and 4 °C for 30 minutes (Eppendorf 5424 R, FA 45 24-11 rotor). The supernatant containing the solubilized protein complexes was then subjected to an ultracentrifugation on top of a 98% (w/v) glycerol cushion at 385,900 x g, 4 °C for 30 minutes (Optima MAX XP centrifuge, Beckman Coulter, Brea, USA; MLA130 rotor, Beckman Coulter, Brea, USA). The resulting supernatant was carefully removed and centrifuged once more at the same ultracentrifugation settings without a glycerol cushion. The resulting, magnetosome-free supernatant was then concentrated using Amicon Ultra 0.5 mL spin columns with a 3 kDa molecular weight cut-off at 4 °C according to the manufacturer’s protocol to a protein concentration of 3.5 mg mL^-1^ in a volume of 300 µL. 100 µL of this solution were then separated on a Superose 6 Increase 10/300 GL with a flow rate of 0.3 mL per minute on an Äkta pure system (GE Healthcare, Uppsala, Sweden). Chromatography was performed in buffer E containing 50 mM NaCl, 0.005% (w/v) LMNG and 0.5 mL fraction were collected. 15 µL were then used for analysis by SDS-PAGE and Western blotting.

#### Carbonate extraction of magnetosome proteins

Carbonate extractions were performed according to a modified protocol described previously (Clements et al., 2009). For carbonate extraction of membrane associated MM proteins, 400 µL of magnetosomes with an OD_492_ of 6 in buffer E (corresponding to 0.28 µg mL^-1^ protein) were pelleted by centrifugation (20 min, 21,000 x g, 10 °C). The supernatant was carefully discarded, and the pellet was resuspended in 400 μL 100 mM NaCO_3_ (pH 11.3) by vortexing. Samples containing 400 µL buffer E or 400 µL of 1% SDS in buffer E, served as negative and positive controls, respectively. After incubation for 1.5 h at RT on a roll mixer, samples were pelleted by centrifugation (20 min, 21,000 x g, 10 °C; Eppendorf 5424 R, FA-45-24-11 rotor). The supernatants were transferred to ultracentrifuge tubes, refilled with respective buffers (ad 600 µL) and centrifuged (30 minutes, 385,900 x g, 4 °C; Optima MAX XP centrifuge, Beckman Coulter, Brea, USA; MLA130 rotor, Beckman Coulter, Brea, USA) twice. While the magnetosome pellets were resuspended in the same buffers again, the supernatant was neutralized and precipitated in 150 μL aliquots by addition of 1.5 mL ice-cold 90% (v/v) acetone, 10% (v/v) trichloroacetic acid (TCA), 10 mM NaCl, and incubation at −20 °C overnight. Supernatant samples were then centrifuged at 21,000 x g 4 °C for 20 min and the supernatant was discarded. Precipitates were washed with 1 mL ice-cold acetone and pelleted by centrifugation (21,000 x g, 4 °C for 20 min) three times. Pellets were then dried at RT to remove residual acetone, resuspended in 40 µL buffer E and 5x SDS loading buffer with short incubation in a sonication bath, and stored at 20 °C until further use. Resuspended magnetosomes were washed in buffer E three times before magnetosome pellets were dissolved in 300 µL buffer E and 5x SDS loading buffer and stored at 20 °C until use. 20 µL aliquots were used for SDS-PAGE and subsequent Western blot analyses.

Further extractions were performed with buffer E and basic (0.1 M N-cyclohexyl-3-aminopropanesulfonic acid (CAPS-NaOH), pH 11), acidic (0.1 M glycine-HCl, pH 2.5), or high salt (10 mM Hepes, 1 mM EDTA, 1 M NaCl, pH 7.4) buffer solutions under the same conditions, except that extraction was performed for 20 h with buffer exchanges after 1, 3, and 16 h.

#### Co-immunoprecipitation

For co-immunoprecipitation experiments, 400 µL of magnetosomes with an OD_492_ of 6 in buffer E (corresponding to 0.28 µg mL^-1^ protein) were pelleted by centrifugation (15 min, 21,000 x g, 10 °C). The supernatant was carefully discarded, and the pellet was resuspended in 100 μL buffer 1 (10 mM HEPES, 1 mM EDTA, 50 mM NaCl, 0.1 mM PMSF, 0.5% (v/v) Triton X-100, pH 7.4). After incubation for 12 h at 4 °C on a roll mixer, samples were pelleted by centrifugation (15 min, 21,000 x g, 10 °C). Subsequently, 100 µL of the supernatant were transferred to new 1.5 mL reaction tube and 400 μL buffer 2 (10 mM HEPES, 1 mM EDTA, 50 mM NaCl, 0.1 mM PMSF, 0.1% (v/v) Triton X-100, pH 7.4) were added (input). Prior to immunoprecipitation, 25 μL of the GFP-Trap® Agarose (Chromotek, Planegg-Martinsried, Germany) affinity resin were equilibrated three times with 500 μL buffer 2, sedimented by centrifugation (5 min, 2,500 x g, 4 °C) and the supernatant was discarded. 450 μL of the diluted input were added to the equilibrated beads and after incubation for 1 h at 4 °C on a roll mixer, beads were sedimented by centrifugation (5 min, 2,500 x g, 4 °C; Eppendorf 5424 R, FA-45-24-11 rotor). The supernatant (unbound) and 50 µL of the unused input were diluted 1:1 with 2x SDS-sample buffer (100 mM Tris, 20% (v/v) glycerol (v/v), 4% (w/v) SDS, 0.2% (w/v) bromophenol blue, 200 mM DTT, pH 6.8) and stored at −20°C until further analysis. The beads were washed three times with 500 μL buffer 2, sedimented by centrifugation (5 min, 2,500 x g, 4 °C) and the supernatant was discarded. Finally, the beads were transferred to new 1.5 mL reaction tubes, resuspended in 80 μL 2x SDS-sample buffer, boiled for 5 min at 95 °C to dissociate immunocomplexes (IP) from beads and sedimented by centrifugation (5 min, 2,500 x g, 10 °C). Samples were then evaluated via SDS-PAGE and subsequent Western immunoblots by loading 10 µL (input) and 20 µL (unbound, Co-IP) aliquots, respectively.

#### Sample preparation for mass spectrometry

Magnetosome samples were prepared for mass spectrometry using filter aided sample preparation (FASP) as described elsewhere (Wiśniewski, 2016). Briefly, aliquots containing 100 µg of total protein were reduced with tris(2-carboxyethyl)phosphine followed by mixing with 200 µL of buffer UA (8 M urea in 0.1 M Tris/HCl, pH 8.5) and loading on a Microcon YM 30 (Merck-Millipore, Darmstadt, Germany) filtration device by centrifugation. Subsequently, proteins were alkylated by adding iodoacetamide in buffer UA and digested for 18 hours at 37 °C with trypsin at a ratio of 1:100 (trypsin:protein) in 40 µL 50 mM Tris/HCl. Peptides were eluted by centrifugation with 100 µL 50 mM Tris/HCl, twice. Eluates were pooled and resulting peptides were purified by Pierce C18 Tips 100 µL (Thermo Fisher Scientific, Waltham, USA). Therefore, C18 tips were wetted with 200 µL of 70% acetonitrile and equilibrated with 200 µL 3% acetonitrile. Peptides were bound by aspirating and dispensing ten times. Bound peptides were washed and eluted with water and 60% acetonitrile, respectively. Eluted peptides were dried and stored at −80 °C until further use.

#### Mass spectrometry of magnetosomes

Purified peptides were reconstituted with 10 µL 0.1% acetic acid and analyzed by reversed phase liquid chromatography (LC) electrospray ionization (ESI) MS/MS using a QExactive Hybrid-Quadrupol-Orbitrap mass spectrometer (Thermo Fisher Scientific, Waltham, USA). In brief, nano reversed phase LC columns (20 cm length x 100 µm diameter) packed with 3.0 µm C18 particles (Dr. Maisch GmbH, Ammerbuch Entringen, Germany) were used to separate the purified peptides with an EASY nLC 1000 system (Thermo Fisher Scientific, Waltham, USA). The peptides were loaded with buffer A (0.1% acetic acid) and subsequently eluted by a non-linear gradient of 166 min from 2% to 99% buffer B (0.1% acetic acid, 99.9% acetonitrile) at a flow rate of 300 nL min^-1^. A full scan was recorded in the Orbitrap with a resolution of 70,000. The twelve most abundant precursor ions were consecutively isolated by the quadrupole and fragmented via higher-energy collisional dissociation (HCD) with a normalized collision energy of 27.5%. MS2 scans were recorded with a resolution of 17,500. Unassigned charge states, singly charged ions, as well as ions of charge 7 and higher were rejected and the lock mass correction was enabled.

Database searching and quantification was done with MaxQuant version 1.6.3.4. (Cox and Mann, 2008) with the published genome sequence of MSR (GenBank: CP027526.1) (Uebe et al., 2018b). The MaxQuant generic contaminants database was used. Database search was based on a strict tryptic digestion with two missed cleavages permitted. Carbamidomethylating on cysteine was considered as a fixed modification and oxidation of methionine as a variable modification. MaxQuant computed LFQ intensities were loaded into Perseus 1.6.2.2 (Tyanova et al., 2016) and log2 transformed. Putative contaminants, reverse hits, and proteins identified by site only were removed and a list containing proteins identified in all samples was exported to Excel. Mean log2 differences and respective P values were obtained by a two-sided two sample t-test over three biological replicates.

#### Epifluorescence microscopy

For epifluorescence microscopy, MSR strains were grown in 3 mL FSM at 2% O2 at 28 °C without shaking for approximately 20 h. When required, gene expression was induced by addition of 100 ng mL^-1^ anhydrotetracycline or 2 mM isopropyl β-D-1-thiogalactopyranoside (IPTG). For imaging, cells were immobilized on 1% agarose pads supplemented with FSM media components (except for peptone and yeast extracts). Therefore, 3 µL of cell suspension were pipetted on agarose pads and covered with a coverslip. Samples were then imaged with an Olympus BX81 microscope equipped with a 100× UPLSAPO100XO objective (NA1.4), an OrcaER camera (Hamamatsu), and DIC contrast. Epifluorescence micrographs were recorded in Z-stacks with 750 ms exposure time per image and then deconvoluted employing 200 iterations of the Richardson-Lucy algorithm (Richardson, 1972; Lucy, 1974) using the DeconvolutionLab 2.0.0 plugin (Sage et al., 2017) the ImageJ Fiji package (Schindelin et al., 2012).

#### Structured illumination microscopy

3D-SIM (striped illumination at 3 angles and 5 phases) was performed on an Eclipse Ti2-E N-SIM E fluorescence microscope (Nikon) equipped with a CFI SR Apo TIRF AC 100×H NA1.49 Oil objective lens, a hardware based ‘perfect focus system’ (Nikon), LU-N3-SIM laser unit (488/561/640 nm wavelength lasers) (Nikon), and an Orca Flash4.0 LT Plus 17 sCMOS camera (Hamamatsu). Calibration of the objective correction collar and SIM grating focus was performed with TetraSpeck fluorescent beads (T-7279 TetraSpeck microspheres). For 3D-SIM imaging, cells were prepared as described above except that 8-Well µ-Slides with 1.5H (170±5 µm) D 263 M Schott glass bottom (ibidi GmbH, Gräfelfing, Germany) were used. 3D SIM z-series were acquired with 120 nm z-step spacing and exposure times in the range of 20 to 100 ms at 25 to 75% laser power. EM515/30 and EM595/31 filters and fluorescence excitation with 488 nm and 561 nm lasers were used for imaging of GFP and mCherry/mScarlet-I, respectively. Image reconstruction was performed in NIS-Elements 5.01 (Nikon) using the ‘stack reconstruction’ algorithm with the following parameter settings: The ‘illumination modulation contrast’ was set to ‘auto’. The ‘high resolution noise suppression’ was set to 0.1.

#### Transmission electron microscopy

For transmission electron microscopy (TEM) analysis, cells were grown at 28 °C under microaerobic conditions (2% O2) over-night. One milliliter cell suspension (OD565 ∼ 0.2-0.3) was then concentrated by centrifugation at 1.000 x g for 3 min, followed by resuspension in ∼50 µL of residual medium. Afterwards cells were adsorbed onto carbon coated copper mesh grids (CF200-CU, Electron Microscopy Sciences, Pennsylvania) and washed with ddH_2_O twice. Images were recorded using Zeiss EM 902A and Jeol JEM-1400 Plus electron microscopes at an accelerating voltage of 80 kV. For data processing, interpretation, and analysis, the software packages DigitalMicrograph (Gatan) and the ImageJ Fiji package (Schindelin et al., 2012) were used. For determinations of magnetite particle numbers per cell, at least 57 cells were analyzed and at least 300 particles were measured for analysis of magnetite particle diameters.

Magnetosome chain formation was analyzed from TEM images as depicted in Figure S2. Briefly, TEM images were segmented using the ImageJ plugin Trainable Weka Segmentation v3.2.28 (Arganda-Carreras et al., 2017) for extraction of cell boundaries and a difference of Gaussians (DoG) method (sigma1 = 1, sigma2 = 2) (Bundy, 1986) to enhance the edges of the images and enable extraction of magnetosomes by thresholding. After manual curation to prevent erroneous segmentation of polyphosphate granules, binary magnetosome images were subjected to particle number analyses using the inbuilt “Analyze Particles” function of Fiji. For neighbor analyses, the “Neighbor Analysis” function of the Fiji BioVoxxel toolbox plugin 2.5.0 (J. Brocher, 2015) was used in particle neighborhood mode with 35 nm neighborhood radius. Intracellular magnetosome distribution maps were generated with MicrobeJ (Ducret et al., 2016) from at least 593 particles and 16 cells.

#### Bioinformatic analyses

For the detection of remote MFP homologies, Hidden Markov Model-based HHPred analyses were performed (Zimmermann et al., 2018). Using individual MFP sequences or MFP alignments as queries, the domain of unknown function DUF4870 (HOTT) and the Tic20 protein families were consistently the only hits with significant statistical support in various databases like PFAM (Mistry et al., 2021), TIGRFAMs (Haft et al., 2001), or COG (Galperin et al., 2021). Subsequently, 5200 protein sequences of the identified DUF4870 (HOTT) and Tic20 protein families were retrieved from the InterPro database (Blum et al., 2021). After an initial Cluster analysis of sequences (CLANS) (Frickey and Lupas, 2004), 229 DUF4870 (HOTT) and Tic20 family proteins of bacterial, archaeal and eukaryotic origin were selected to achieve a broad phylogenetic distribution. These sequences were again analyzed by CLANS using default settings for 150,000 iterations. Only if the P value for a pair of sequences is less than 10^−5^ in the all-against-all BLAST search, the corresponding edges between nodes are shown as gray or black lines. To generate maximum-likelihood trees of MamF-like proteins or DUF4870 (HOTT) and TIC20 family proteins 59 sequences were aligned using MAFFT 7.474 (Katoh et al., 2019), respectively. Trimmed alignments (TrimAI 1.3 Capella-Gutiérrez et al., 2009, no gaps and gap threshold 0.7, respectively) were then used to infer maximum-likelihood trees with IQ-Tree 1.6.11 (Trifinopoulos et al., 2016) under the LG+G4 or mtInv+F+I+G4 models as suggested by ModelFinder (Kalyaanamoorthy et al., 2017). Bootstrap support was derived by ultrafast bootstrap approximation with 1000 iterations. The phylogenetic trees were visualized and annotated by iTOL online tool (Letunic and Bork, 2019).

To infer the phylogenetic distribution of the DUF4870 (HOTT) and Tic20 families, a total of 16840 rRNA sequences (16S or 18S) from organisms encoding these protein families were initially retrieved from the NCBI nucleotide database. After filtering for sequences of at least 1200 or 1655 nt length and removal of sequences with similarities above 97% (CD-Hit, Huang et al., 2010), a total of 456 (HOTT) and 297 (Tic20) rRNA sequences were aligned using MAFFT 7.490 (Katoh et al., 2019), respectively. Trimmed alignments (TrimAI 1.3, no gaps) were then used to infer maximum-likelihood trees with IQ-Tree 1.6.12 (Trifinopoulos et al., 2016) under the GTR+F+R10 or TIM3+F+R3 models as suggested by ModelFinder (Kalyaanamoorthy et al., 2017). Bootstrap support was derived by ultrafast bootstrap approximation with 1000 iterations. The phylogenetic trees were visualized and annotated by iTOL online tool (Letunic and Bork, 2019). All sequences and alignments were edited and analyzed using Geneious 8.1.4 (Biomatters, Auckland, New Zealand).

All sequences and alignments were edited and analyzed using Geneious 8.1.4 (Biomatters, Auckland, New Zealand).

For protein signal sequence predictions, SignalP 5.0 (Almagro Armenteros et al., 2019) was used. Hydrophobicity analyses were performed using the grand average of hydropathy (GRAVY) calculator (http://www.gravy-calculator.de) (Kyte and Doolittle, 1982; Peschke et al., 2019).

Figures and RMSD of superimpositions of AlphaFold database (AF) predictions (Jumper et al., 2021; Varadi et al., 2022) of Tic20/HOTT superfamily members with the PDB (Protein data bank) structure of Tic20 (*C. reinhardtii*, PDB ID: 7xZI) were generated with PYMOL (The PyMOL Molecular Graphics System, Version 2.0 Schrödinger, LLC).

#### Quantification and statistical analysis

All statistical analyses were performed with GraphPad Prism 7 software (GraphPad Software, Inc., La Jolla, CA, USA). All data were analyzed using two-tailed Student’s t or Mann-Whitney U tests, respectively. A P value of less than 0.05 was considered statistically significant. Further information about statistical details and methods is indicated in the figure legends, text, or methods. If not stated otherwise, values are given as mean ± standard deviation of the indicated sample size. Violin plots, bar plots, and intracellular magnetosome distribution maps were generated by Fit-o-Mat 0.752 (Möglich, 2018), Prism 7 software (GraphPad) and MicrobeJ (Ducret et al., 2016), respectively. Unless indicated otherwise, all experiments were performed at least twice.

### Supplementary Text

#### Altered composition of the MM in MFP mutants

In addition to the eleven proteins that showed a significantly decreased abundance in MM fractions of strain ΔF3, our quantitative proteomic analyses also revealed that 26 proteins are significantly enriched compared to the WT (data table S4). Among these proteins, the three MAPs MamC, MamW, and MamX showed a three-fold increase (Fig. 3D). While increased MamC levels could be confirmed by Western immunoblots (fig. S5A), the deletion of *mamC* or *mamW* had only very weak effects on magnetosome formation (Scheffel et al., 2008; Lohße et al., 2011). Since MamX, on the other hand, plays a role in magnetosome redox control (Raschdorf et al., 2013), the enrichment of these MAPs does not seem to play a role for the ΔF3 phenotype. Beside these MAPs, 23 significantly enriched proteins have no known function in magnetosome biomineralization and are encoded outside the genomic magnetosome island that contains almost all MAP genes (Lohße et al., 2011). Strikingly, 13 of these proteins contain predicted secretory N-terminal signal sequences (SignalP 5.0) (data table S4) (Almagro Armenteros et al., 2019) and thus likely reside within the periplasm or the outer membrane. To test for a putative role in magnetosome biogenesis, we fluorescently labeled three of the most strongly enriched proteins (fig. S5, B and C). When produced in the WT, all proteins showed a peripheral cell localization, whereas mCherry-labeled MmeA, a MAP with a known N-terminal signal peptide (Richter et al., 2007), showed a magnetosome chain-like localization pattern. Thus, at least the tested proteins (MSR1_01270, MSR1_09970, MSR1_21580) do not localize to the MM *in vivo.* We thus concluded that only proteins with a decreased abundance in ΔF3 MM extracts are relevant for our analyses.

**Fig. S1.**
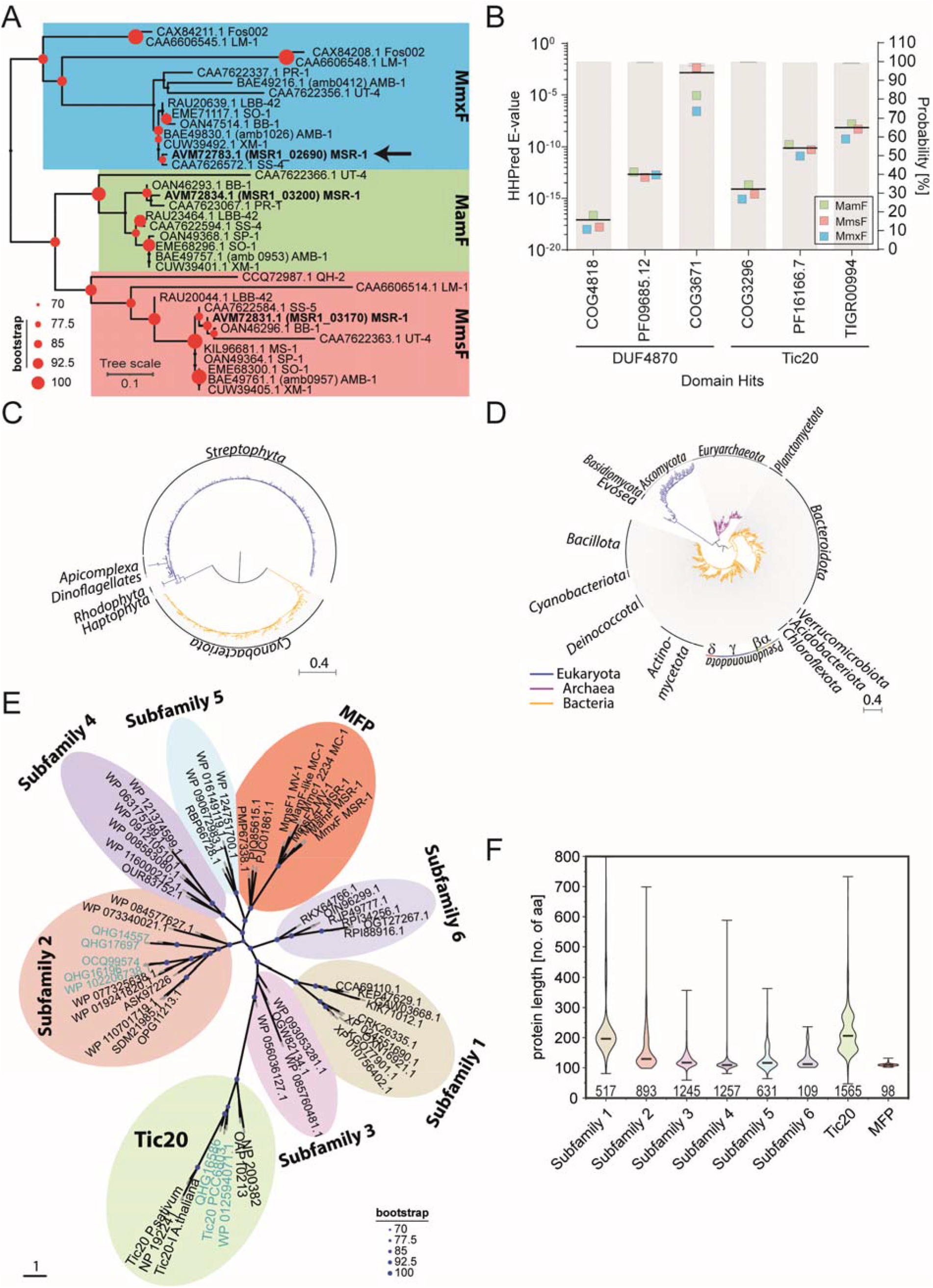
MFPs are members of a common Tic20/HOTT protein superfamily, which are distributed in all domains of life. **(A)** A phylogenetic analysis of various MFPs reveals that one of the newly identified MAPs (MSR1_02690) is an ortholog of MmxF (amb1026) from *M. magneticum* AMB-1 (Rawlings et al., 2014). The tree is based on 36 MFPs sequences from different magnetotactic bacteria inferred under the best-fitting LG+G4 substitution model. Grouped MmxF, MamF and MmsF homologs are shaded with blue, green, and red boxes, respectively. MSR MFPs are highlighted in bold and MmxF is marked with an arrow. Red dots above branches are bootstrap values in percent, denoted in the legend. Scale bar represents expected substitutions per site. Accession numbers are listed in data table S1. **(B)** HHPred analysis of the MSR MFPs reveals significant homologies to the DUF4870 (HOTT) and Tic20 protein families. E-values (average indicated by a black line) indicate highly significant homologies of MamF, MmsF, and MmxF (green, red, blue squares, respectively) to different HOTT and Tic20 protein family models. The averaged hit probability of the templates is represented by grey bars with grey lines indicating the standard deviation. Only hits with E-values < than 10^-3^ are shown. **(C)** A phylogenetic analysis reveals a narrow distribution of the Tic20 family. The phylogenetic tree is based on 16S/18S rRNA sequences from 297 organisms containing at least one Tic20 family protein. The tree was inferred under the best-fitting TIM3+F+R3 substitution model. Scale bar represents expected substitutions per site. **(D)** A phylogenetic analysis reveals the wide distribution of the HOTT family in all domains of life. The phylogenetic tree is based on 16S/18S rRNA sequences from 456 organisms containing at least one HOTT family protein. The tree was inferred under the best-fitting GTR+F+R10 substitution model. Scale bar represents expected substitutions per site. **(E)** Unrooted phylogenetic tree of the Tic20/HOTT superfamily inferred under the best-fitting mtInv+F+I+G4 substitution model. The MFPs and six other HOTT groups form distinct subfamilies within the HOTT family that is widely separated from the TIC20 family. HOTT Subfamily 1 exclusively includes eukaryotic sequences whereas groups 2–6 contain only prokaryotic representatives. Cyanobacteria (in teal) are the only organisms containing HOTT and Tic20 family proteins. Bootstrap values (blue dots) are denoted in the legend. Scale bar represents expected substitutions per site. Accession numbers are listed in data table S1. **(F)** Comparison of protein length distributions (in aa) in the Tic20/HOTT superfamily using violin plots. Please note the short length of most HOTT family members in comparison to Tic20. The min, max, and mean values are indicated by bars. The number of analyzed proteins of each group [N] is given below. (See also Figure 1 and data table S1)

**Fig. S2.**
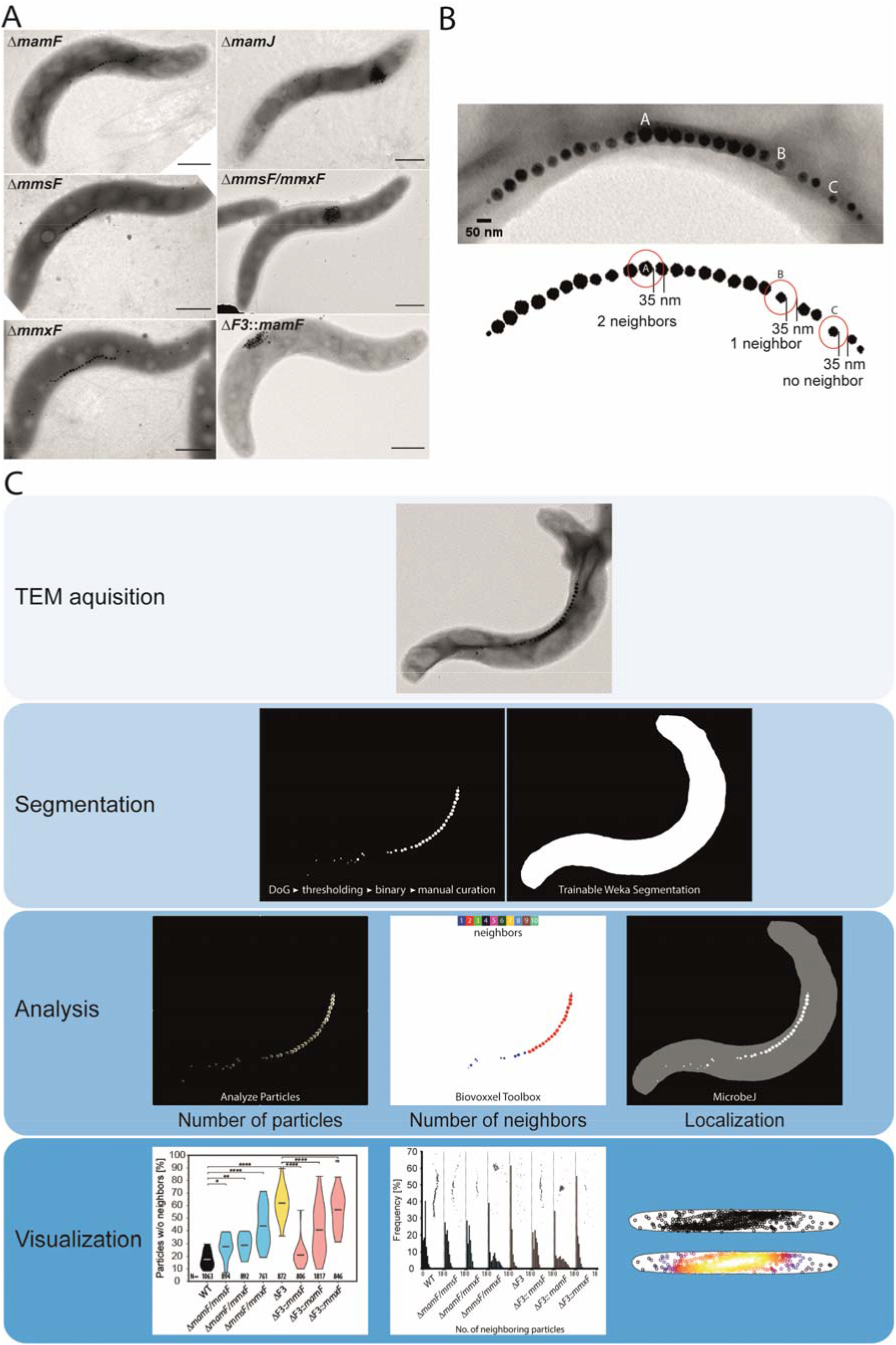
Characterization of MSR deletion phenotypes using quantitative magnetosome neighbor analysis. **(A)** Overview of different MSR deletion phenotypes. Representative TEM micrographs of MFP single deletions show WT-like phenotypes (left), while mutants lacking *mmsF* and *mmxF* but contain *mamF* phenocopy the Δ*mamJ* strain (right) (Scale bars: 500 nm). **(B)** MSR WT magnetosomes with an average size of 35 nm are arranged in almost perfect chain-like structures in which most particles have exactly two neighboring particles within a distance of 35 nm (see particle A). Exceptions can be found at the end of chains where a wider particle spacing can be observed (e.g. particle B that has only one neighboring particle within 35 nm) or particle C that has no neighbor. In the absence of MCs the frequency of neighbor-less particles increases strongly while strains with aggregated magnetosomes frequently have neighbor numbers ≥ 3. **(C)** For qMNA images of different mutant strains were collected by TEM. Micrographs were subsequently segmented into magnetosomes and cell bodies. Segmented binary images were then used to analyze the overall number of magnetosomes per cell, the number of neighboring magnetosomes of each individual particle and the distribution of intracellular magnetosome localizations using Fiji plugins. Visualization of the results was performed using Fit-o-mat 0.752 (Möglich, 2018), Prism 7 (GraphPad Prism Software Inc., San Diego, California), and MicrobeJ (Ducret et al., 2016). (See also Figure 2, data table S3)

**Fig. S3.**
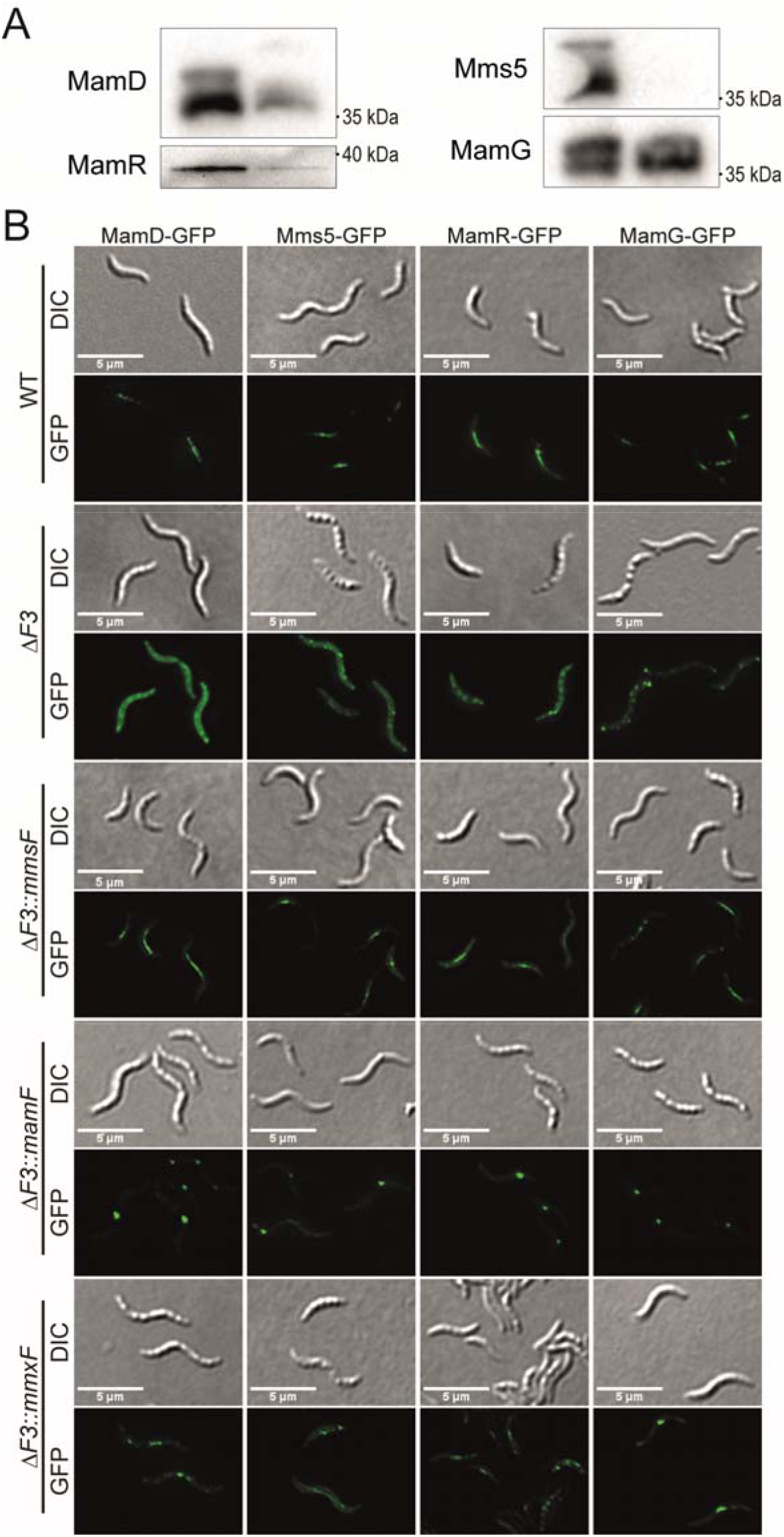
Verification of the ΔF3 proteome analysis. (A) Western blot analyses (αGFP) of MM fractions from WT and ΔF3 strains expressing *mamG*-, *mamD*-, *mms5*-, and *mamR-GFP* confirm a strong reduction of MamD, Mms5, and MamR in ΔF3 magnetosomes compared to the WT. In contrast, MamG-GFP is equally abundant in both strains. (B) Epifluorescence imaging of WT, ΔF3, and complemented ΔF3 strains producing MamD, Mms5-, MamR-, or MamG-GFP fusion proteins reveals MM targeting of MamD-, Mms5-, and MamR-GFP by all three MFPs. In ΔF3, MamD-, MamR-, and partially Mms5-GFP show only a soluble localization whereas MamG-GFP exclusively localizes in a punctuate pattern. DIC, differential interference contrast image. (Scale bars: 5 µm) (See also Fig. 3)

**Fig. S4.**
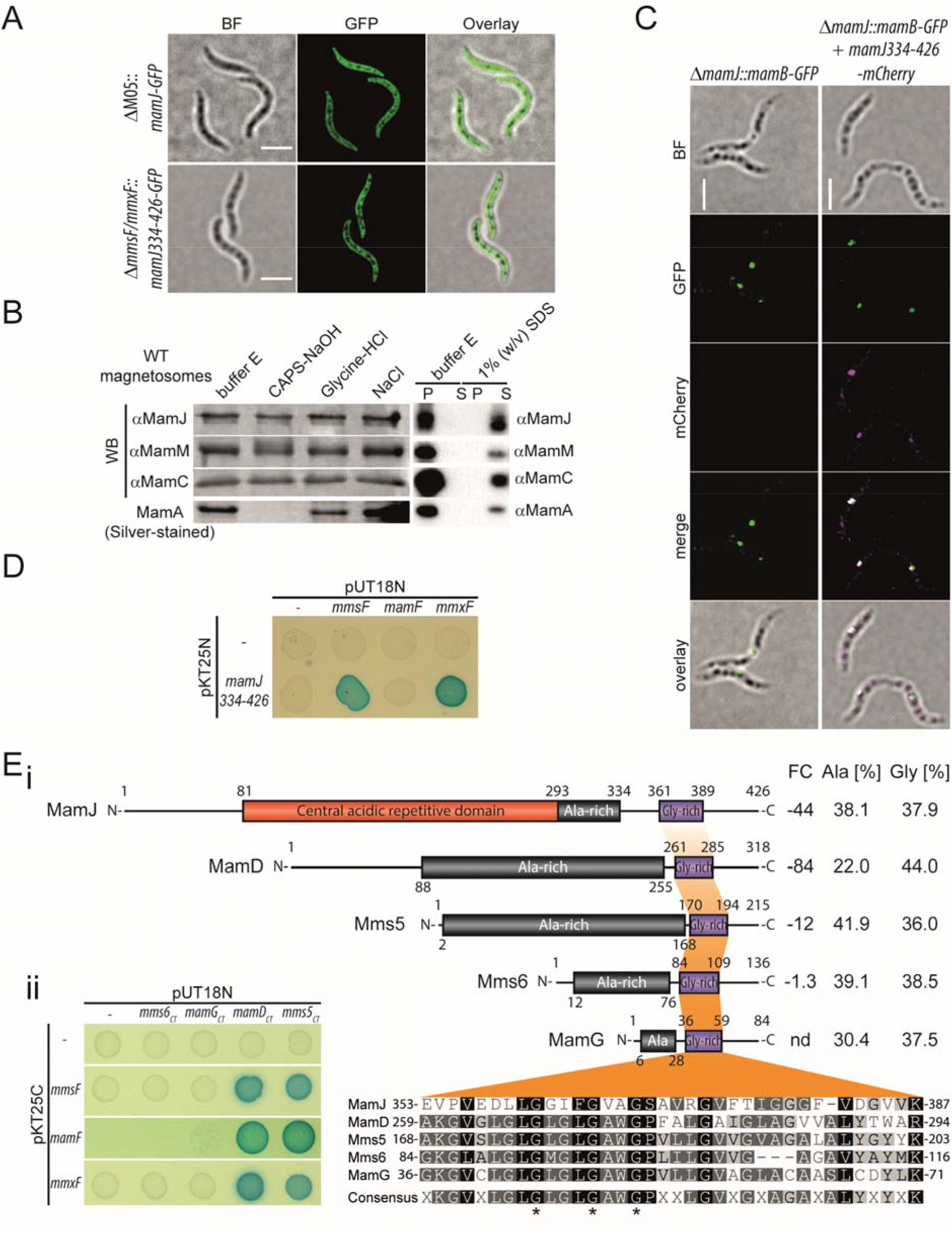
Characterization of potential MFP substrate proteins. **(A)** 3D-Structured illumination microscopy of MamJ-GFP reveals a soluble localization pattern in the MSR ΔM05 mutant that lacks all magnetosome genes. Similarly, MamJ334-426-GFP has a soluble localization in the Δ*mmsF*/*mmxF* mutant that forms Δ*mamJ*-like magnetosome clusters but is unable to target MamJ to the MM. BF, bright field image. (Scale bars: 2 µm) **(B)** MamJ is resistant to prolonged treatments (20 h) with basic (0.1 M N-cyclohexyl-3-aminopropanesulfonic acid (CAPS), pH 11), acidic (0.1 M glycine, pH 2.5), or high salt (1 M NaCl, 10 mM Hepes, 1 mM EDTA, pH7.4) buffer solutions. Magnetosome pellet fractions were loaded on SDS-PAGE and subsequently analyzed by Western blot (αMamJ, αMamM, αMamC) or silver staining (MamA). In an independent experiment, magnetosomes were incubated for one hour in 1% SDS as a positive control. The sample was fractionated into supernatant and magnetosome pellets by centrifugation. TCA-precipitated supernatant and the pellet fractions were loaded onto SDS-PAGE. Subsequent Western blot immunodetections of MamJ, MamM, and MamC indicate an integral membrane protein behavior for all three proteins as they could be only extracted by SDS. Contrarily, the MM associated protein MamA could be readily extracted by SDS and CAPS-buffer treatments. Buffer E treatments served as negative controls. **(C)** 3D-Structured illumination colocalization microscopy of MamB-GFP and MamJ334-426-mCherry in ΔmamJ indicate that residues 334-426 mediate MM targeting of MamJ (see also Fig. 3, Fig. 4, fig. S2A and Supplementary text). Micrographs are maximum intensity projections. MamB-GFP is colored in green; MamJ-mCherry is colored in magenta; White-colored regions in the overlay indicate colocalization. BF, bright field image. (Scale bars: 2 µm) **(D)** Interaction analysis of MamJ using a BACTH assay based on reconstitution of CyaA adenylate cyclase activity in the cya− strain E. coli BTH101. Fusion of proteins to the fragmented catalytic CyaA domains T25 and T18 from Bordetella pertussis only confers cAMP-dependent expression of a lacZ reporter gene upon protein-protein interaction-mediated reconstitution of CyaA. lacZ expression is indicated by blue color formation and enhanced growth on X-Gal containing M63 maltose-mineral salts agar. The MamJ C-terminal domain interacts with MmsF and MmxF but not MamF in BACTH assays. **(E**i) Domain structures of putative MFP substrates MamJ, MamD and Mms5 as well as the MamD-homologs MamG and Mms6 which are not targeted to the MM in an MFP-dependent manner (fold change (FC) of the proteomic analysis is indicated; nd, not detected). Alanine and glycine contents of the alanine-(black) and glycine-rich (purple) domains are given in %, respectively. Numbers above and below the proteins represent domain boundaries or protein lengths, respectively. Multiple aa sequence alignment of the glycine-rich domains. Amino acid residues are shaded using the Rasmol color scheme. A consensus motif is given below the alignment. The numbers at the beginning and end of each line indicate the position of the first and last amino acid of the respective protein within the alignment. Asterisks indicate position of conserved glycin residues. **(Eii)** Interaction analysis of MamD, Mms5, MamG, Mms6 C-terminal domains using a BACTH assay based on reconstitution of CyaA adenylate cyclase activity in the *cya^−^* strain E. coli BTH101. BACTH assay showing the interaction between the C-terminal domains of MamD and Mms5 with MamF, MmsF, and MmxF. Note that the highly homologous proteins MamG and Mms6 do not interact with the MFPs. (See also Fig. 4)

**Fig. S5.**
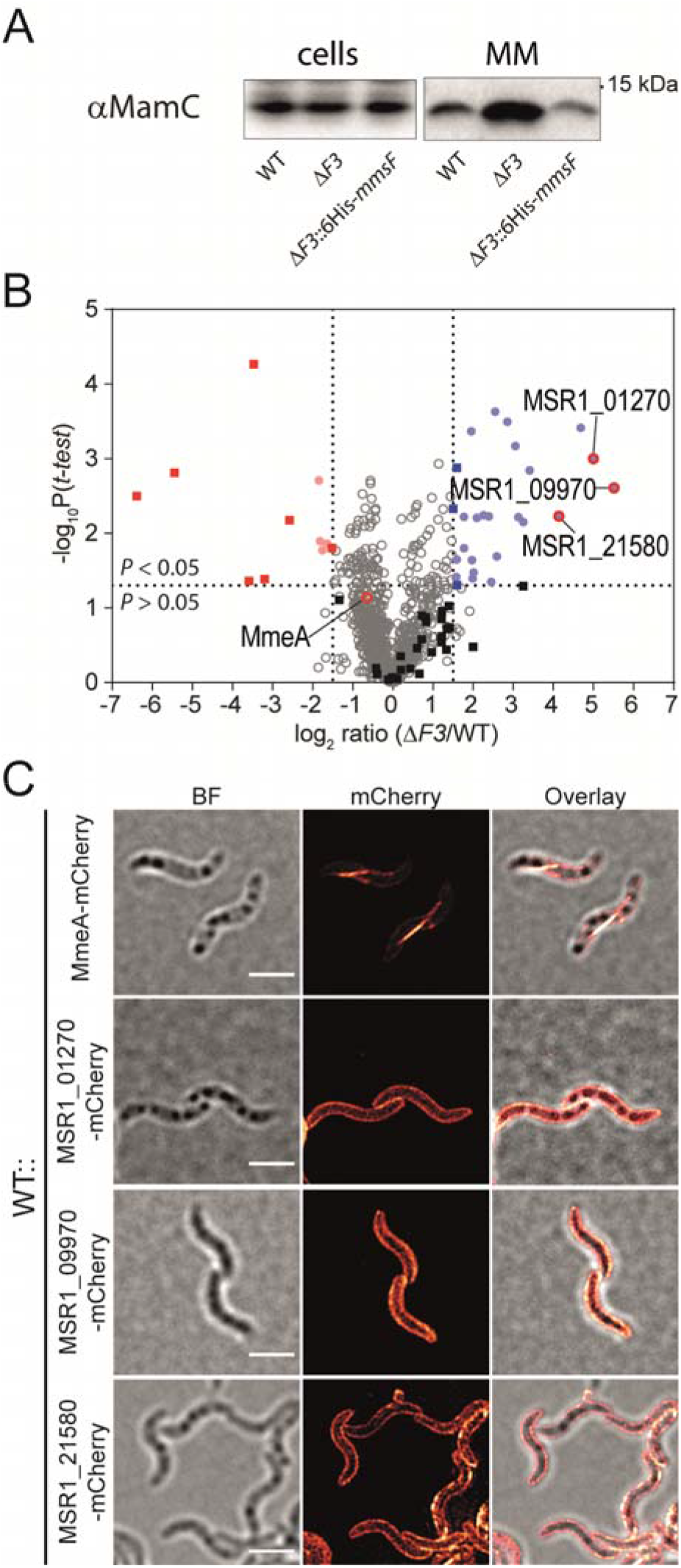
Verification of the proteomic data. **(A)** Volcano plot showing the log2-fold depletion (red) or enrichment (blue) of proteins in purified magnetosome fractions of ΔF3 compared to the WT as determined by LC-MS/MS. Circles represent non-MAI encoded proteins, non-significant proteins are depicted by grey or black symbols, respectively. Proteins with predicted N-terminal signal peptide sequences and high enrichment in ΔF3 used for mCherry-labelling are highlighted (red circles). MmeA, whose abundance remained unaffected in ΔF3 MM fractions and used as a reference for mCherry labelling is also highlighted. Coloring otherwise identical to Fig. 3D. (See also Data table S3) **(B)** 3D-Structured illumination microscopy confirms that mCherry-labeled proteins (see Figure S3D) with potential secretory signal sequences are localized near the cell surface in the MSR WT. Except for the known MAP MmeA no protein showed MC localization. BF, bright field image. (Scale bars: 2 μM) (See also Figure 3)

**Table S1.** Strains used in this study.

| Strain | Genotype/Description | Reference/Source |
| --- | --- | --- |
| ***E. coli*** |  |  |
| DH5α | Host for cloning:<br>F<sup>-</sup> Φ80*lacZ*ΔM15 Δ(*lacZYA-argF*) U169 *recA1*<br>*endA1 hsdR17* (*r<sub>k</sub><sup>-</sup>*, *m<sub>k</sub><sup>+</sup>*) *phoA supE44 thi-1*<br>*gyrA96 relA1 λ<sup>-</sup>* | Invitrogen |
| WM3064 | Donor strain for conjugation:<br>*thrB1004 pro thi rpsL hsdS lacZ*ΔM15 RP4-1360<br>Δ(*araBAD*)567 Δ*dapA*1341::[*erm pir* (wt)] | William Metcalf at UIUC |
| BTH101 | BACTH assay reporter strain:<br>F<sup>-</sup> *cya-99 araD139 galE15 galk16 rpsL1* (Str<sup>R</sup>)<br>*hsdR2 mcrA1 mcrB1* | Euromedex |
| ***M. gryphiswaldense*** |  |  |
| Wild type (WT) | MSR-1 R3/S1; Rif<sup>R</sup>, Sm<sup>R</sup> spontaneous mutant | (Schultheiss et al., 2004) |
| WT::*mamD-GFP* | WT with *mamD-GFP* after Tn5 insertion, Tet<sup>R</sup> | This study |
| WT::*mamR-GFP* | WT with *mamR-GFP* after Tn5 insertion, Tet<sup>R</sup> | This study |
| WT::*mms5-GFP* | WT with *mms5-GFP* after Tn5 insertion, Tet<sup>R</sup> | This study |
| WT::*mamG-GFP* | WT with *mamG-GFP* after Tn5 insertion, Tet<sup>R</sup> | This study |
| WT::P<sub>tet</sub>-*mmeA-mCherry* | WT with *mmeA-mCherry*<br>after Tn5 insertion, Kan<sup>R</sup> | This study |
| WT::*MSR1_01270-mCherry* | WT with *MSR1_01270-mCherry*<br>after Tn7 insertion, Cam<sup>R</sup> | This study |
| WT::*MSR1_09970-mCherry* | WT with *MSR1_09970-mCherry*<br>after Tn7 insertion, Cam<sup>R</sup> | This study |
| WT::*MSR1_21580-mCherry* | WT with *MSR1_21580-mCherry*<br>after Tn7 insertion, Cam<sup>R</sup> | This study |
| WT::*MSR1_03900-mCherry* | WT with *MSR1_03900-mCherry*<br>after Tn7 insertion, Cam<sup>R</sup> | This study |
| ΔM05 | Non-magnetic mutant with large MAI operon deletions | (Dziuba et al., 2020) |
| ΔM05::*mamJ-GFP* | ΔM05 with *mamJ-GFP*<br>after Tn5 insertion, Tet<sup>R</sup> | This study |
| Δ*mamJ* | *mamJ* deletion strain | (Scheffel et al., 2006) |
| Δ*mamJ*::*mamJ-GFP* | Δ*mamJ* with *mamJ-GFP*<br>after Tn5 insertion, Tet<sup>R</sup> | This study |
| Δ*mamJ*::*mamJ1-80-GFP* | Δ*mamJ* with *mamJ1-80-GFP*<br>after Tn5 insertion, Tet<sup>R</sup> | This study |
| Δ*mamJ*::*mamJ81-333-GFP* | Δ*mamJ* with *mamJ81-333-GFP*<br>after Tn5 insertion, Tet<sup>R</sup> | This study |
| Δ*mamJ*::*mamJ334-426-GFP* | Δ*mamJ* with *mamJ334-426-GFP*<br>after Tn5 insertion, Tet<sup>R</sup> | This study |
| Δ*mamJ*::*mamJΔ335-343-GFP* | Δ*mamJ* with *mamJΔ334-343-GFP*<br>after Tn5 insertion, Tet<sup>R</sup> | This study |
| Δ*mamJ*::*mamJΔ335-348-GFP* | Δ*mamJ* with *mamJΔ334-348-GFP*<br>after Tn5 insertion, Tet<sup>R</sup> | This study |
| Δ*mamJ*::*mamJΔ335-353-GFP* | Δ*mamJ* with *mamJΔ334-353-GFP*<br>after Tn5 insertion, Tet<sup>R</sup> | This study |
| Δ*mamJ*::*mamJΔ335-358-GFP* | Δ*mamJ* with *mamJΔ334-358-GFP*<br>after Tn5 insertion, Tet<sup>R</sup> | This study |
| Δ*mamJ*::*mamJΔ378-426-GFP* | Δ*mamJ* with *mamJΔ378-426-GFP*<br>after Tn5 insertion, Tet<sup>R</sup> | This study |
| *ΔmamJ::mamJΔ396-426-GFP* | *ΔmamJ* with *mamJΔ396-426-GFP* after Tn5 insertion, Tet<sup>R</sup> | This study |
| *ΔmamJ::mamJΔ407-426-GFP* | *ΔmamJ* with *mamJΔ407-426-GFP* after Tn5 insertion, Tet<sup>R</sup> | This study |
| *ΔmamJ::mamJΔ417-426-GFP* | *ΔmamJ* with *mamJΔ417-426-GFP* after Tn5 insertion, Tet<sup>R</sup> | This study |
| *ΔmamJ::mamB-GFP* | *ΔmamJ* with chromosomally encoded *mamB-GFP*, expressed from the native *mamB* locus | This study |
| *ΔmamJ::mamB-GFP::P<sub>lac</sub>-mamJ334-426-mCherry* | *ΔmamJ* with chromosomally encoded *mamB-GFP*, expressed from the native *mamB* locus and *mamJ334-426-mCherry* inserted into the Tn7 locus | This study |
| *ΔmamF* | *mamF* deletion strain | (Lohße et al., 2014) |
| *ΔmmsF* | *mmsF* deletion strain | (Lohße et al., 2014) |
| *ΔmmxF* | *mmxF* deletion strain | This study |
| *ΔmamF/mmsF* | *mamF*, *mmsF* double deletion strain | (Lohße et al., 2014) |
| *ΔmamF/mmxF* | *mamF*, *mmxF* double deletion strain | This study |
| *ΔmmsF/mmxF* | *mmsF*, *mmxF* double deletion strain | This study |
| *ΔmmsF/mmxF::mamJ334-426-GFP* | *ΔmmsF/mmxF* *mamJ334-426* after Tn5 insertion, Tet<sup>R</sup> |  |
| *ΔF3* | *mamF*, *mmsF*, *mmxF* triple deletion strain | This study |
| *ΔF3::mamF* | Tn5 mediated complementation of *ΔF3* with *mamF*, Tet<sup>R</sup> | This study |
| *ΔF3::mmsF* | Tn5 mediated complementation of *ΔF3* with *mmsF*, Tet<sup>R</sup> | This study |
| *ΔF3::mmxF* | Tn5 mediated complementation of *ΔF3* with *mmxF*, Tet<sup>R</sup> | This study |
| *ΔF3::6His-mmsF* | *ΔF3* with *6His-mmsF* after Tn5 insertion, Kan<sup>R</sup> | This study |
| *ΔF3::P<sub>tet</sub>-mCherry-mamK* | *ΔF3* with inducible *mCherry-mamK* after Tn5 insertion, Kan<sup>R</sup> | This study |
| *ΔF3::P<sub>tet</sub>-mScarlet-I-mamY* | *ΔF3* with inducible *mScarlet-I-mamY* after Tn5 insertion, Kan<sup>R</sup> | This study |
| *ΔF3ΔmamJ::mamJ-GFP* | *ΔF3ΔmamJ* with *mamJ-GFP* after Tn5 insertion, Tet<sup>R</sup> | This study |
| *ΔF3::GFP-mamF* | *ΔF3* with *GFP-mamF* after Tn5 insertion, Tet<sup>R</sup> | This study |
| *ΔF3::GFP-mmsF* | *ΔF3* with *GFP-mmsF* after Tn5 insertion, Tet<sup>R</sup> | This study |
| *ΔF3::mamD-GFP* | *ΔF3* with *mamD-GFP* after Tn5 insertion, Tet<sup>R</sup> | This study |
| *ΔF3::mamR-GFP* | *ΔF3* with *mamR-GFP* after Tn5 insertion, Tet<sup>R</sup> | This study |
| *ΔF3::mms5-GFP* | *ΔF3* with *mms5-GFP* after Tn5 insertion, Tet<sup>R</sup> | This study |
| *ΔF3::mamG-GFP* | *ΔF3* with *mamG-GFP* after Tn5 insertion, Tet<sup>R</sup> | This study |
| *ΔF3::mamD-GFP::mmsF* | *ΔF3* with *mamD-GFP* and *mmsF* after Tn5 insertion, Kan<sup>R</sup>, Tet<sup>R</sup> | This study |
| *ΔF3::mamR-GFP::mmsF* | *ΔF3* with *mamR-GFP* and *mmsF* after Tn5 insertion, Kan<sup>R</sup>, Tet<sup>R</sup> | This study |
| *ΔF3::mms5-GFP::mmsF* | *ΔF3* with *mms5-GFP* and *mmsF* after Tn5 insertion, Kan<sup>R</sup>, Tet<sup>R</sup> | This study |
| *ΔF3::mamG-GFP::mmsF* | *ΔF3* with *mamG-GFP* and *mmsF* after Tn5 insertion, Kan<sup>R</sup>, Tet<sup>R</sup> | This study |
| *ΔF3::mamD-GFP::mamF* | *ΔF3* with *mamD-GFP* and *mamF* after Tn5 insertion, Kan<sup>R</sup>, Tet<sup>R</sup> | This study |
| *ΔF3::mamR-GFP::mamF* | *ΔF3* with *mamR-GFP* and *mamF* after Tn5 insertion, Kan<sup>R</sup>, Tet<sup>R</sup> | This study |
| $\Delta F3::mms5\text{-GFP}::mamF$ | $\Delta F3$ with *mms5-GFP* and *mamF* after Tn5 insertion, Kan<sup>R</sup>, Tet<sup>R</sup> | This study |
| $\Delta F3::mamG\text{-GFP}::mamF$ | $\Delta F3$ with *mamG-GFP* and *mamF* after Tn5 insertion, Kan<sup>R</sup>, Tet<sup>R</sup> | This study |
| $\Delta F3::mamD\text{-GFP}::mmxF$ | $\Delta F3$ with *mamD-GFP* and *mmxF* after Tn5 insertion, Kan<sup>R</sup>, Tet<sup>R</sup> | This study |
| $\Delta F3::mamR\text{-GFP}::mmxF$ | $\Delta F3$ with *mamR-GFP* and *mmxF* after Tn5 insertion, Kan<sup>R</sup>, Tet<sup>R</sup> | This study |
| $\Delta F3::mms5\text{-GFP}::mmxF$ | $\Delta F3$ with *mms5-GFP* and *mmxF* after Tn5 insertion, Kan<sup>R</sup>, Tet<sup>R</sup> | This study |
| $\Delta F3::mamG\text{-GFP}::mmxF$ | $\Delta F3$ with *mamG-GFP* and *mmxF* after Tn5 insertion, Kan<sup>R</sup>, Tet<sup>R</sup> | This study |
| $\Delta F3::mmsF^{D34N, D36N, D37N, E38N}$ | $\Delta F3$ with *mmsF*<sup>D34N, D36N, D37N, E38N</sup> after Tn5 insertion, Tet<sup>R</sup> | This study |
| $\Delta F3::mmsF^{D34N, R35Q, D36N, D37N, E38N}$ | $\Delta F3$ with *mmsF*<sup>D34N, R35Q, D36N, D37N, E38N</sup> after Tn5 insertion, Tet<sup>R</sup> | This study |
| $\Delta mamKY$ | *mamK*, *mamY* double deletion strain | (Toro-Nahuelpan et al., 2019) |
| $\Delta F3\Delta mms5\Delta mamK\Delta mamY::GFP\text{-}mmsF$ | *mamF*, *mmsF*, *mmxF*, *mms5*, *mamK*, *mamY* codeletion strain; $\Delta F3\Delta mms5\Delta mamK\Delta mamY$ with *GFP-mmsF* after Tn5 insertion, Tet<sup>R</sup> | This study |
| $\Delta A13\Delta mmxF\Delta mms5$ | *mamGFDC*, *mms5*, *mms6*, *mamXY* codeletion strain | (Zwiener et al., 2021) |
| $\Delta A13\Delta mmxF\Delta mms5\Delta mamR$ | *mamGFDC*, *mms5*, *mms6*, *mamXY*, *MamR* codeletion strain | This study |
| $\Delta A13\Delta mmxF\Delta mms5\Delta mamR::mmsF$ | $\Delta A13\Delta mmxF\Delta mms5\Delta mamR$ with *mmsF* after Tn5 insertion, Tet<sup>R</sup> | This study |

**Table S2.**
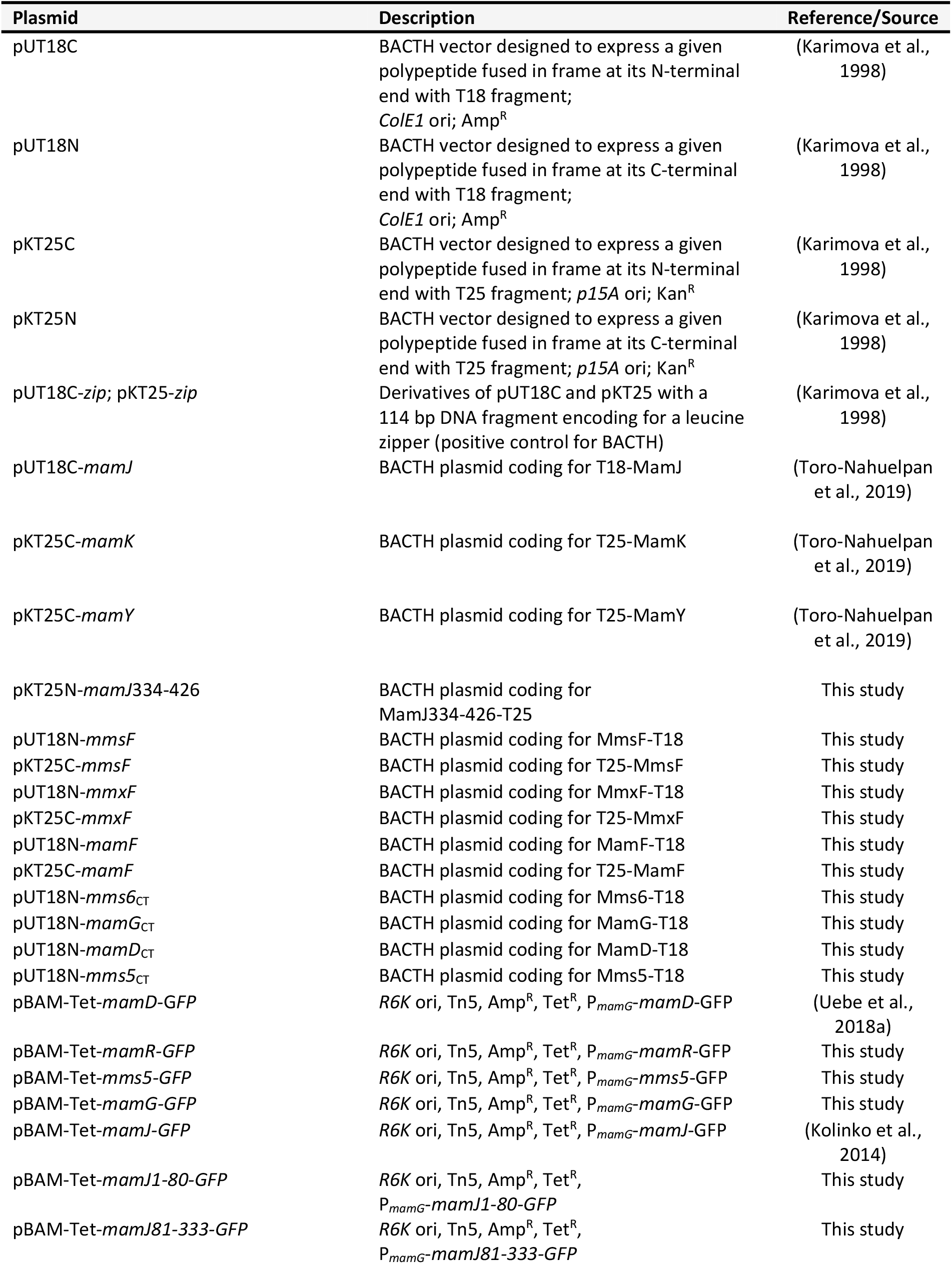
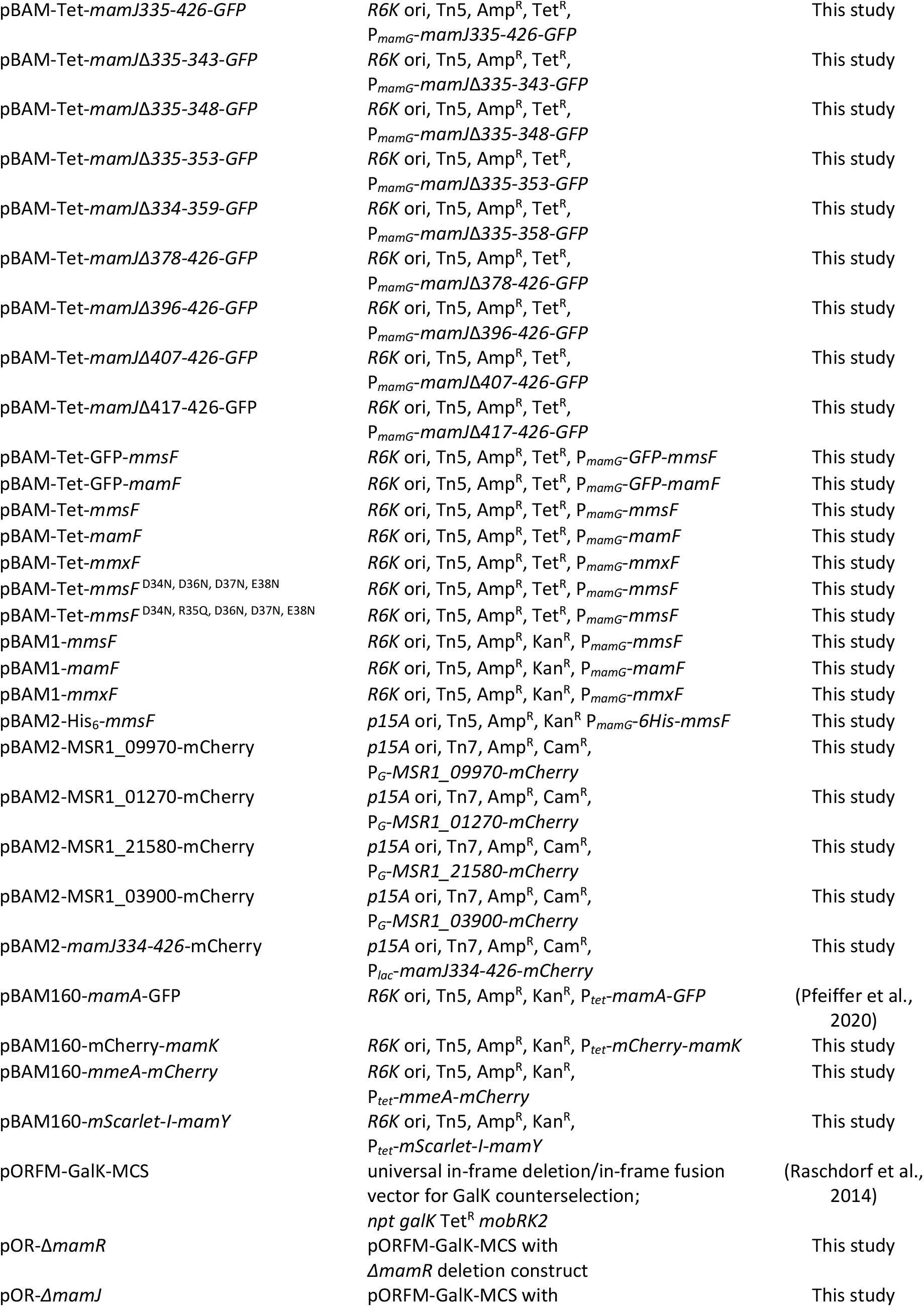
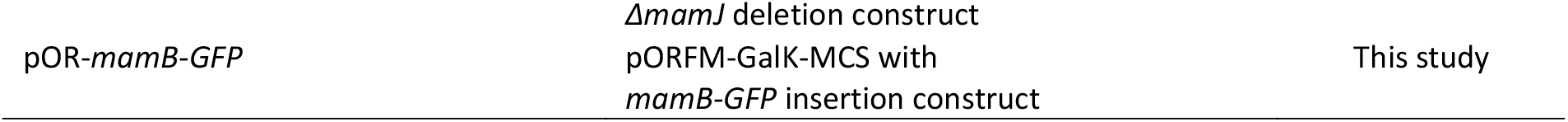
Plasmids used in this work.

**Table S3.**
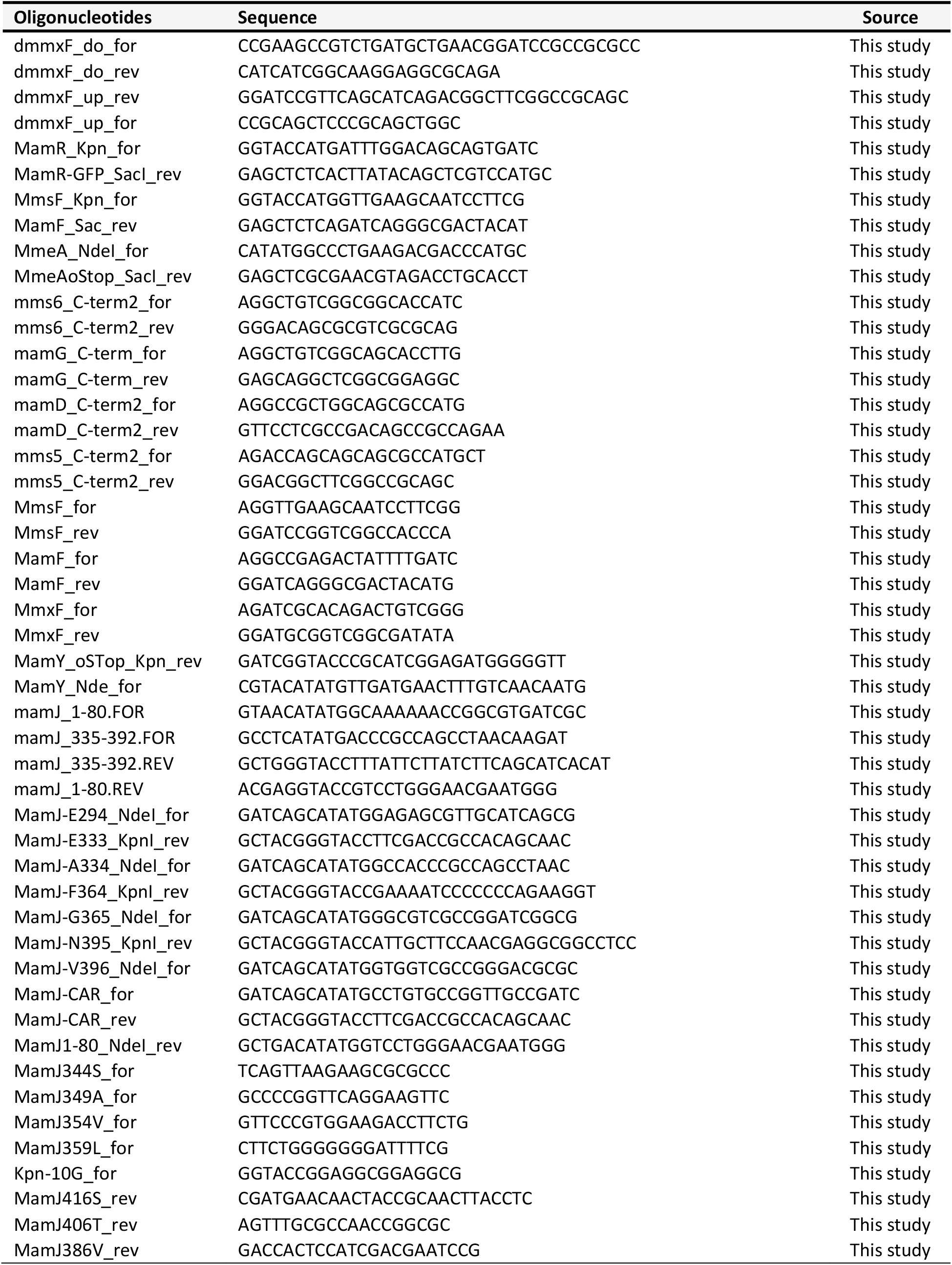
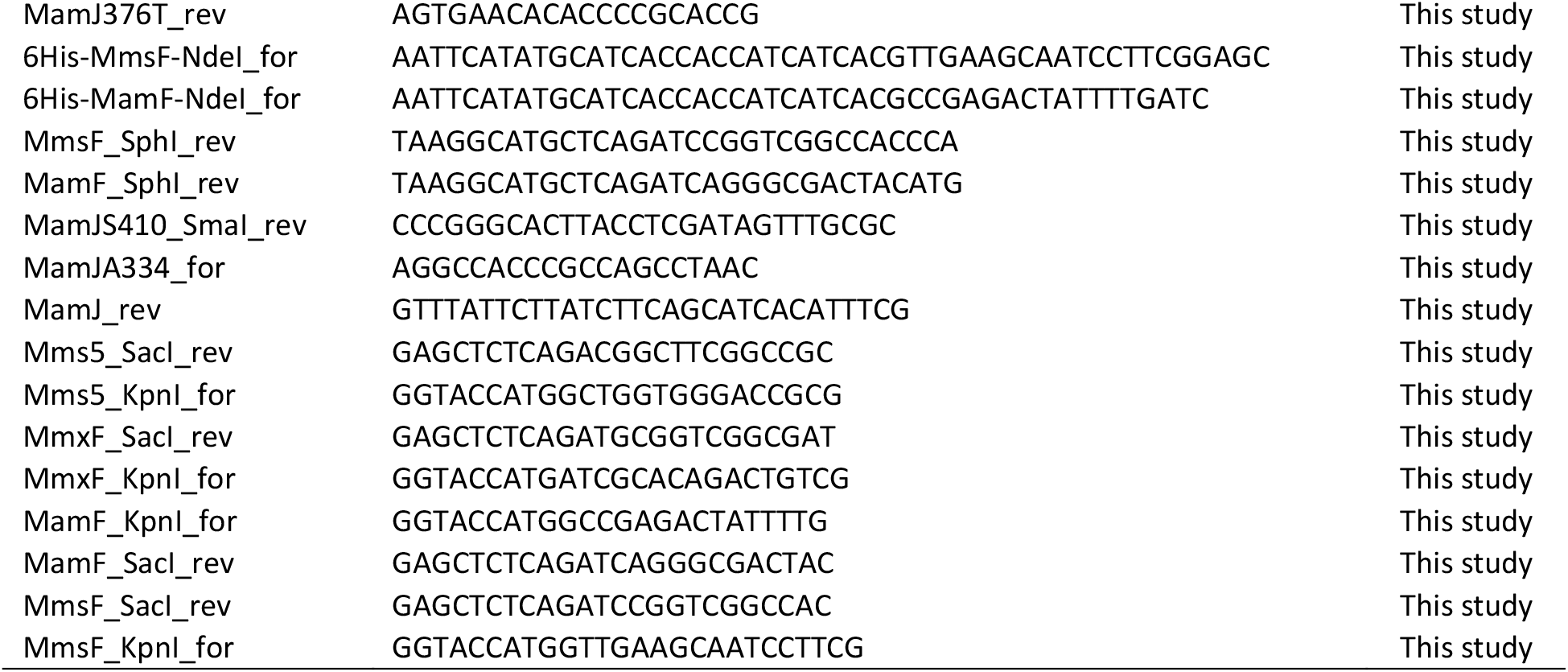
PCR Primers used in this work.

#### Data tables (separate excel files)

Data table S1. Accession numbers of proteins used for phylogenetic analyses of MFPs (Figure S1A) and the whole Tic20/HOTT superfamilies (Figure S1E)

Data table S2. Accession numbers of proteins used for CLANS analysis (Figure 1A) and structural/multiple sequence alignments (Figure 1 B and C)

Data table S3. Data for magnetosome crystal size (Figure 2A), magnetosome number per cell (Figure 2B), qMNA (Figure 2D and E), and magnetotaxis analyses (Figure 2F).

Data table S4. List of proteins enriched or depleted in the ΔF3 MM fraction as determined by quantitative proteomic analysis. Related to Figures 3D and S3D.

Data table S5. Data for magnetosome crystal size analysis for Figure 5B.

## Notes

### Competing Interest Statement

The authors have declared no competing interest.

